# Exploring the Microbial Geobiological Pattern Across the Serpentinization Sites through Metagenomic and Elemental Composition Analyses

**DOI:** 10.1101/2025.09.26.678891

**Authors:** Triparna Mukherjee, Anupkumar Rai, Ian C Marshall, Rudra Prasad Mondal, Punit Kohli, Shimin liu, Satya Harplani, Divya Prakash

## Abstract

It is well-known that ultramafic rocks can continuously generate hydrogen through the serpentinization process associated with dynamic geochemical and geobiological interactions. This study aims to gain an improved understanding of these dynamic processes through the understanding of diverse microbial populations in order to maximize the geologic hydrogen production potential. The water samples were collected from a near surface serpentinite site in Northern California, USA. Elemental analyses of these water samples revealed the presence of essential dissolved minerals such as magnesium, calcium, and potassium, along with trace elements including iron, cobalt, and nickel. These elements act as cofactors for enzymes involved in microbial metabolic processes and Non-metric Multidimensional Scaling analysis supports this corelation. Furthermore, the analysis of total organic carbon (TOC) showed significant levels of organic carbon, suggesting a link with biological carbon cycling processes. Metagenomic analysis uncovered a diverse microbial community of hydrogen-fueled microbial consortia at the sampling sites, encompassing hydrogenogenic and hydrogenotrophic microorganisms. We examine hydrogen-metabolizing communities, including sulfate-reducing bacteria, acetogens, and methanogens, supported by diversity in hydrogenase enzymes across various sampling sites. These observations are corroborated by genomic accession data and abundance profiling of genes associated with acetogenesis, methanogenesis, and carbon monoxide metabolism. These investigations provide new insights into hydrogen-metabolizing microorganisms in northern California and propose frameworks for optimizing hydrogen production through the inhibition of hydrogen metabolism.

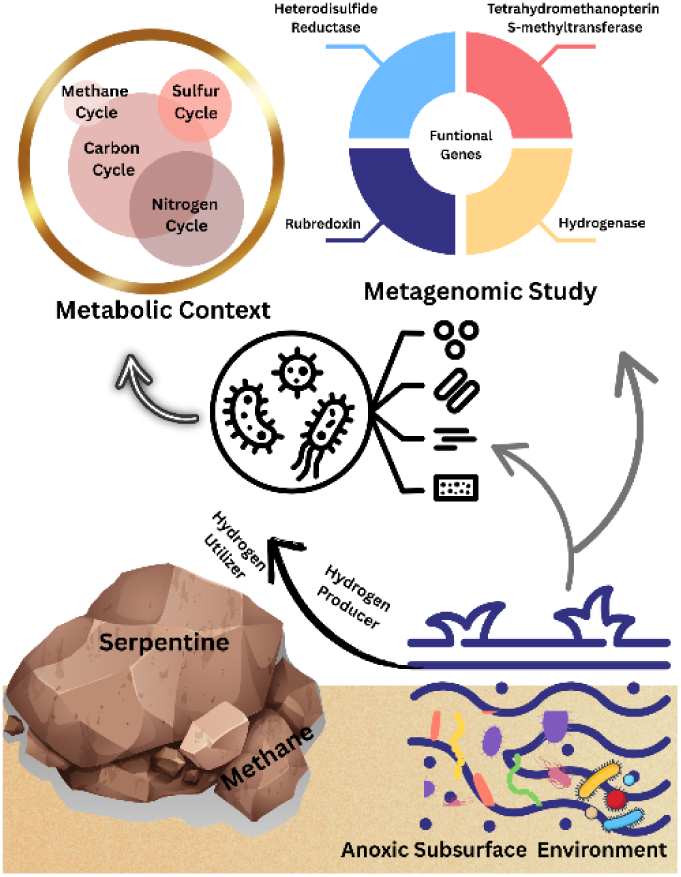

**Highlights:**

- The association of the serpentinization process with dynamic geochemical and geobiological interactions.
- Metagenomic study found an assorted microbial community of hydrogen-fueled consortia at the different sampling sites in Northern California
- Hydrogen-metabolizing communities counted, including sulfate-reducing bacteria, acetogens, and methanogens.

## 1. Introduction

Ultramafic rock, a constituent of the Earth’s mantle, is brought to the surface *via* tectonic uplift. This rock can then undergo serpentinization by interacting with water, releasing hydrogen gas and low-molecular-weight organic compounds(Barbier et al., 2020). Consequently, ultramafic rocks can serve as litho-source rock for hydrogen production *via* the serpentinization process. However, a significant challenge in progressing hydrogen technology through this method is the potential for induced (micro)-seismic activity due to the matrix swelling, as well as the microbiological factors associated with low-temperature reservoirs(Osselin et al., 2022). If not properly managed, these factors can lead to the consumption of the generated H2 and thus reduction of hydrogen production. Microbiology is neglected in the planning and management of underground hydrogen storage. (Bertagni et al., 2022; Dopffel et al., 2023; Gregory et al., 2019; Tremosa et al., 2023). Hydrogen serves as both an electron donor and an energy source for various microbial groups. Beyond the potential for hydrogen loss, which has economic implications, biological processes can trigger other potentially hazardous, costly, and environmentally harmful outcomes, including H2S formation, the production of CH4 and other hydrocarbons, corrosion, and various geochemical reactions ((Bertagni et al., 2022), (Liu and Conrad, 2011; Schwab et al., 2023; Thaysen et al., 2021)). The foremost concerns regarding three hydrogen-utilizing microbial metabolisms are sulfate reduction, methanogenesis, and acetogenesis ((Dopffel et al., 2023),(Schwab et al., 2023; Strobel et al., 2023; Thaysen et al., 2021)). These processes might take place at any hydrogen production and storage facility, but biological complexities currently hinder the development of universally applicable conclusions. Therefore, it is essential to understand the microbial consortia at serpentinization sites to prevent the utilization of hydrogen gas and maximize the hydrogen yield under the reservoir conditions.

A near surface serpentinite geological formation was identified in northern California. It is known for its serpentine bionetwork, where microbes may thrive in an extreme geochemical environment characterized by hyperalkalinity, extremely low redox potential, and a nominal ionic concentration. These geochemical considerations make a discernible environment, influencing microbial associations that vary between the sources of groundwater. The site is known not only for its alkalinity, which promotes the proliferation of specific microbes, but also for the extreme conditions in which the plants exist (Morrill et al., 2013). Consequently, this serpentinite site offers an opportunity for further exploration of its geochemical and geobiological aspects in relation to serpentinization processes. This study enhances the scope of such investigations by adapting the comprehensive chemical and microbial composition of this site. To investigate the potential microbial inhabitants of the site, five locationas— designated as Site 1, Site 2, Site 3, Site 4, and Site 5—were chosen for water sampling. This study involved a thorough geochemical investigation in conjunction with a geobiological study. Unlike earlier studies of serpentinization sites in the region, this research offers more detailed insights into geobiological aspects and their link to microbial proliferation. (Morrill et al., 2013). The geobiological profiling includes elemental analysis, physicochemical parameters, metagenomic analysis, and fluorescence and electron microscopy studies of different sampling sites. To explore the microbial interaction patterns considering the elements and associated neighbouring microbes of serpentinization sites, we conducted a genome-based metagenomic study. Previous analysis revolved around the phyla of Parcubacteria and Chloroflexi. Our effort concentrated on hydrogen-utilizing microbes, including hydrogen producers and hydrogenotrophic microbes. Our findings also concentrate on the hydrogen-fuelled microbial consortia present at the sampling site, which include hydrogenogenic and hydrogenotrophic microbes. It involves the investigation of the hydrogen-metabolizing microbial community, categorized as sulfate reducers, acetogens, and methanogens, which supports microbial functionality through the prevalence and diversity of the hydrogenase enzyme across the different sites. These findings are validated by genomic accession and abundance profiling, which also include other genes associated with acetogenesis, methanogenesis, and carbon monoxide metabolism. These studies provide new profiles of hydrogen metabolizing microbes in this geological region, as well as provide new frameworks for increasing hydrogen production *via* the inhibition of hydrogen metabolism.

## 2. Methods

### 2.1 Sampling Site Description and Sample Collection

**Water sampling: -** To ensure the integrity of the samples and minimize potential contamination, meticulous sampling protocols were implemented as described (Sackett et al., 2018). Briefly, pre-sterilized 500 mL glass bottles with butyl rubber stoppers were purged and pressurized with high-purity nitrogen gas (N₂) to displace oxygen (O₂) and maintain anaerobic conditions within the headspace. A battery-operated peristaltic pump, equipped with sterile silicone tubing and disposable needles, was used to collect water samples (Fig. 1S). Needles and tubing were replaced between each sampling site to prevent cross-contamination. 400 mL of water was collected into each bottle, leaving a 100 mL headspace to accommodate gas expansion and maintain anaerobic integrity. Two types of samples were collected from each site: natural water as found *in situ* and water stabilized with sodium sulfide and resazurin to maintain strict anaerobic conditions and serve as a redox indicator. Water samples were collected on November 1, 2024, from the following five different sites, called site 1, site 2, site 3, site 4, and site 5(Fig.2S):

A. Site 1 and Site 2; - A spring containing a large amount of deposited calcium carbonate. Site 1 is the groundwater discharging from springs at the lower elevations of the marine sediments of the Franciscan Subduction Complex (FSC).
B. Site 3; - A site located ∼100 meters downstream of Site 1. Although no bubbling was observed here, it is another known spring containing large amounts of calcium carbonate deposits.
C. Site 4 - A smaller pool located adjacent to site 3 was chosen as another sampling site primarily to increase the number of samples taken.
D. Sites 3 and 4 consist of groundwater discharging from springs at higher elevations from the marine sediments of the Franciscan Subduction Complex (FSC) (Morrill et al., 2013).
E. Site 5; - A pool directly adjacent to the Austin Creek water sampling staging area. It is a shallow pool containing a large amount of plant material (presumably algae) that has large amounts of bubbling from underground. The bubbling was presumably hydrogen or methane from either serpentinization or biological activity.

**Figure 1.**
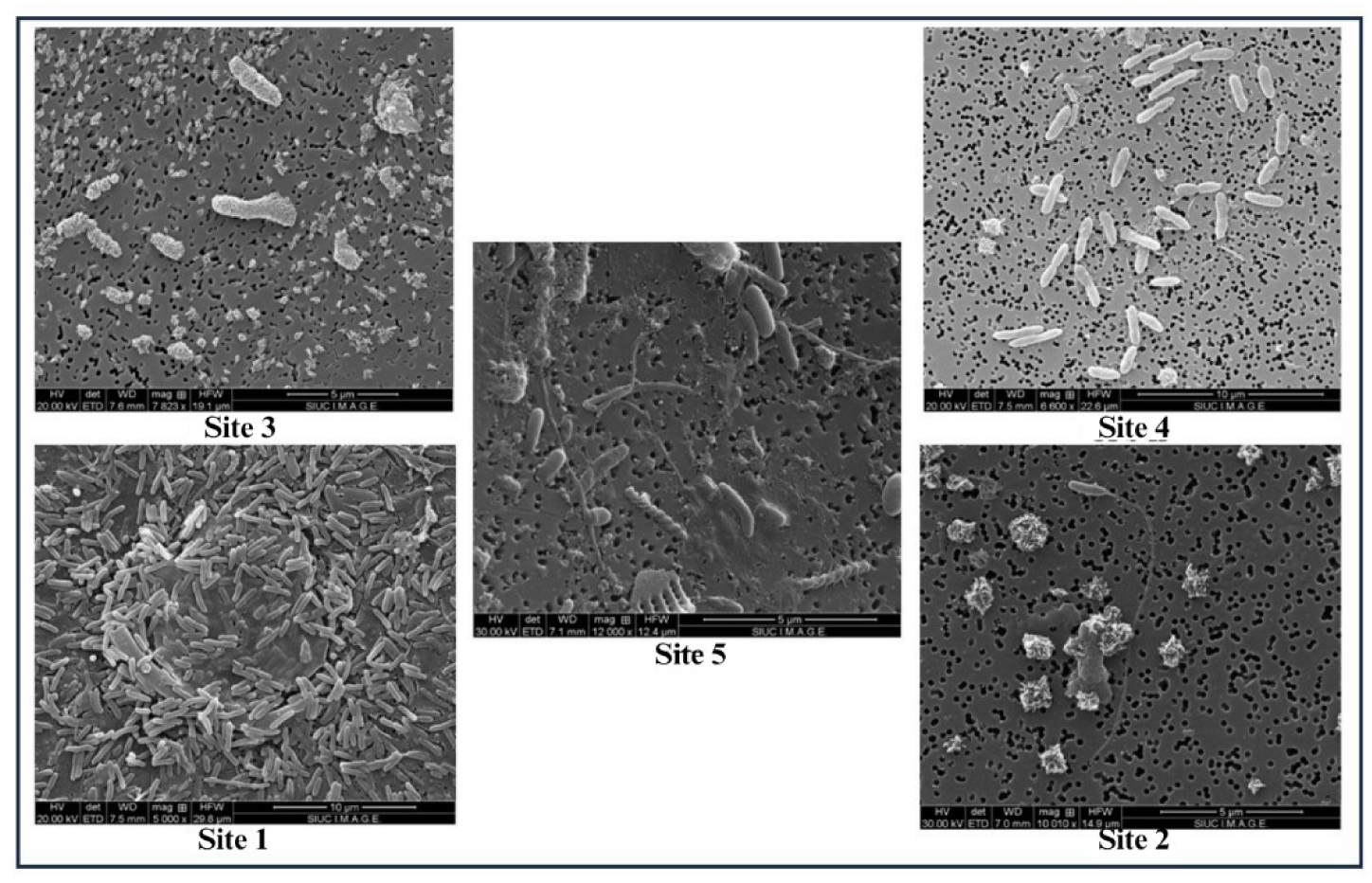
Scanning electron microscopic image of different sampling sites

**Figure 2.**
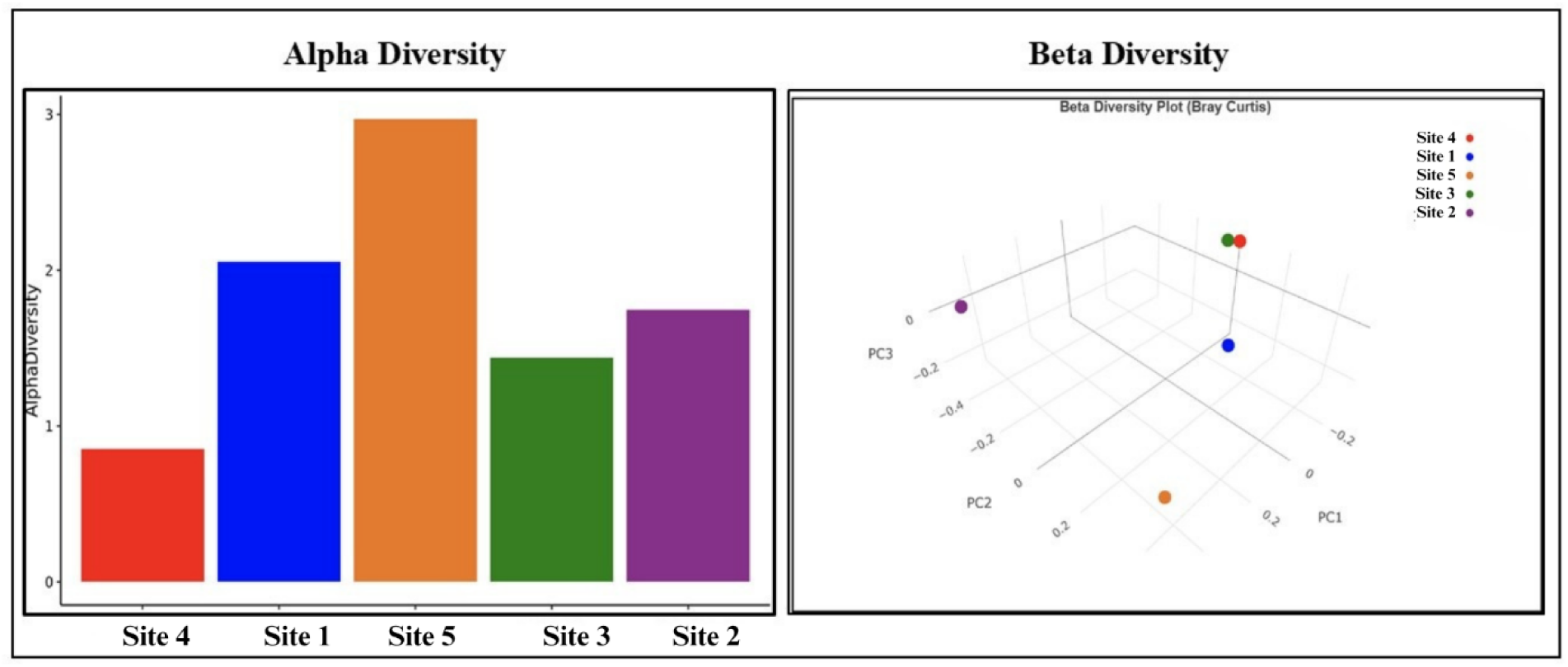
Microbial community divergence pattern across sampling sites a) Alpha diversity, b) Beta diversity; Each dot on the beta diversity plot represents the whole microbial composition profile.

### 2.2 Physicochemical and Geochemical Analyses

Prior to sample collection, temperature (°C) was measured at each sampling site. All samples were immediately placed in an insulated container and transported to the laboratory on ice to minimize microbial activity and maintain sample integrity. The physicochemical parameters were measured on-site using a calibrated dissolved oxygen (DO) probe sensor and a pH meter equipped with a temperature sensor.

### 2.3 Elemental Analysis using inductively coupled plasma mass spectrometry (ICP-MS)

The water samples from different sites were prepared by filtering through 0.22 µm membrane filters to remove particulate matter. Acidification with ultrapure nitric acid was performed to stabilize the samples and ensure accurate measurements of dissolved elements (EPA method 200.8). The ICP-MS instrument was calibrated using a series of multi-element standards, enabling the quantification of trace and major elements.

### 2.4 Total Organic Carbon (TOC), Total Carbon (TC), Total Nitrogen (TN) Measurement

This experiment focused on quantifying total organic carbon (TOC), total carbon (TC), and total nitrogen (TN) in water samples to evaluate organic and inorganic carbon contents and nitrogen levels. For each analysis, 20 mL of the water sample was submitted to the Core Facility for Ecological Analyses at the McLafferty Research Facility, SIU. Samples were centrifuged at 8000 rpm for 30 minutes to remove microbes and cellular debris, which could otherwise interfere with the analysis and result in falsely elevated readings.

The TOC and TC were quantified by measuring the CO₂ released during combustion, while TN was determined based on the nitrogen oxide (NOx) produced. The analysis of sulphate ions was done by a turbidimetric method. This was obtained by adding 3.0 mL 0.1 mol·L-1 BaCl2 solution and exact measured volumes of 0.01 mol·L-1 sulphate solution (0.5 – 3.0 mL) in a 50 mL volumetric flask. The samples were diluted with distilled water, and the turbidity was measured after 5 minutes of standing by 490 nm using a digital spectrophotometer (1 cm glass cell) against distilled water (Anechiţei et al., 2019). The experimental measurement of the turbidity is made at the specific wavelength, which was selected based on the VIS spectrum recorded was for a solution of sulphate ions, with well-known concentration after the addition of BaCl2. It was observed that the BaSO4 precipitate does not have any absorption bands on the entire VIS spectral domain, and this behaviour is characteristic of the white precipitates. Under these conditions, the wavelength that can be used for the turbidity measurements is 490 nm, in agreement with the recommendations from literature (Bîlbă D., 2005). Using Tube test NANOCOLOR Nitrate 50, assay was performed. The reaction uses 2,6-dimethylphenol in a mixture of sulfuric acid and phosphoric acid. Direct nitration of dimethylphenol results in the formation of 4-nitro-2,6-dimethylphenol, depending on the nitrate content of the sample. The absorbance was measured at 350nm.

### 2.5 DNA extraction and sequencing of collected samples

The samples were processed and analyzed with the ZymoBIOMICS® Shotgun Metagenomic Sequencing Service for Microbiome Analysis (Zymo Research, Irvine, CA). DNA extraction was performed using ZymoBIOMICS^TM^ DNA/RNA Miniprep Kit (Cat No: R2002). Genomic DNA samples were profiled with shotgun metagenomic sequencing. Illumina® DNA Library Prep Kit (Illumina, San Diego, CA) with up to 500 ng DNA input following the manufacturer’s protocol using unique dual-index 10 bp barcodes with Nextera® adapters (Illumina, San Diego, CA). All libraries were pooled in equal abundance and the final pool was quantified using qPCR and TapeStation® (Agilent Technologies, Santa Clara, CA). The final library was sequenced on NovaSeq® (Illumina, San Diego, CA). The ZymoBIOMICS® Microbial Community DNA Standard (Zymo Research, Irvine, CA) was used as a positive control for each library preparation. Negative controls (i.e. blank extraction control, blank library preparation control) were included to assess the level of bioburden carried by the wet-lab process.

For the deep sequencing, metagenomic libraries were prepared using the NEBNext Ultra II DNA Library Prep Kit (New England BioLabs, Ipswich, MA). Libraries were quality checked with an Agilent 2100 Bioanalyzer using the DNA High Sensitivity Kit, pooled equimolarly, and gel-purified using a 2% agarose gel and the Qiagen QIAquick Gel Extraction Kit (Qiagen, Germantown, MD, USA). Final pooled libraries were sequenced on an Element AVITI system using 2 × 150 bp paired end reads.

### 2.6 Metagenomic assembly and analysis

Raw sequence reads were trimmed to remove low quality fractions and adapters with Trimmomatic-0.33(Bolger et al., 2014): quality trimming by sliding window with 6 bp window size and a quality cutoff of 20 and reads with size lower than 70 bp were removed. After that, host-derived reads were removed using Kraken2 against some common Eukaryote host genomes(Wood et al., 2019). Low-diversity reads were detected and removed using sdust (https://github.com/lh3/sdust). A read depth of 20 million was covered during this study. The surviving reads were subjected to further taxonomy and functional analyses as follows. Microbial composition was profiled using Sourmash (Irber1, 2016). The GTDB species representative database (RS207) was used for bacterial and archaea identification. Pre-formatted GenBank databases (v. 2022.03) provided by Sourmash (https://sourmash.readthedocs.io/en/latest/databases.html) were also used for virus, protozoa and fungi identification. Reads were mapped back to the genomes identified by Sourmash using BWA-MEM (Li, 2013) and the microbial abundance was determined based on the counts of mapped reads. The resulting taxonomy and abundance information was further analyzed: (1) to perform alpha- and beta-diversity analyses; (2) to create microbial composition barplots with QIIME (Caporaso et al., 2010) (3) to create taxa abundance heatmaps with hierarchical clustering (based on Bray-Curtis dissimilarity); and (4) for biomarker discovery with LEfSe with default settings (p>0.05 and LDA effect size >2)(Segata et al., 2011). Functional profiling was performed using Humann3 including identification of UniRef gene family and MetaCyc metabolic pathways (Beghini et al., 2021). The phylogenetic tree was constructed using GToTree v1.8.4, a command-line tool that powers Hidden Markov Models (HMMs) to find and associate individual copies of marker genes through the genomes. Although GToTree’s default requirement is the presence of a minimum of 50% marker genes, lowering the minimum value to 20% includes MAGs with lower inclusiveness. The resultant phylogenetic tree was further refined using IQ-TREE and visualized with iTOL v7, an interactive tool for exploring and annotating phylogenetic trees.

Another sequencing platform offers a deeper view of microbial profiling, with a depth of up to 60 million reads. In case of deep sequencing Shotgun Metagenomics - Bioinformatic Preprocessing Raw reads were quality-filtered and adapter-trimmed using fastp (Chen, 2023; Chen et al., 2018). Reads were truncated if a sliding window (4 bp) average Phred score dropped below 20. Reads were discarded if the resulting trimmed sequence length was less than 90 bp. Kraken2 was used to assign taxonomic classifications to the remaining sequences with a confidence threshold of 0.2 to reduce spurious assignments (Wood et al., 2019). Sequences were summarized with SeqKit, and host sequences were filtered out again using Kraken2(Shen et al., 2024). KrakenTools was used to extract strain level profiles in report format in combination with custom code to conduct table merging (Lu et al., 2022). Prior to functional annotation remaining, quality filtered microbial reads were dereplicated using VSEARCH (Rognes et al., 2016). Functional annotation was performed with emapper v2.0 using the EggNOG 5.0 database (Cantalapiedra et al., 2021; Huerta-Cepas et al., 2018). KEGG orthologs were used to quantify gene abundances based on the aligned read counts. CCA plot generation was done using Python 3.10.1. Heat map of phylum and species with elements was made by using Python 3.10.1. The co-occurrence heatmap was made by using Python 3.10.1. For deducing the +ve and -ve correlation between microbial population Python 3.10.1 was used.

To get the gene abundance heat map, gene abundance values were log10-transformed after the addition of a pseudocount of 1 to accommodate zero values. A heatmap was generated in R (v4.2) using the heatmap package (v1.0.12). Row labels (KEGG IDs) were simplified by removing redundant EC numbers and taxonomic information for clarity.

### 2.7 Microscopic study

For scanning electron microscopy (SEM), water samples of about 40 ml from five different sites were passed through 0.22µm filter and processed on the filter. 2.5% Glutaraldehyde in 0.1M Phosphate buffer (pH 7.2) with 0.02% Triton X-100 was used as fixative and samples were kept undisturbed for 20 mins at room temp. It was followed by washing of samples with 0.1M Phosphate buffer 3 x 5 minutes. Subsequently samples were washed individually with graded alcohol like 25%, 50%, 70%, 85%, 95%, 3x 100% EtOH 5 min each and dried with CPD. Finally, samples were sputter-coated with 10nm Au/Pd for analysis. The scanning electron microscope was operated under 20-30 kv accelerating voltage maintain the magnification range between 5000x to 10000x.

## 3. Results and Discussion

### 3.1 Physiochemical parameters of formation water from different sites

The ICP-MS analysis revealed (Table 1) the presence of biologically essential elements across all sampling sites, offering insights into biological functions, environmental conditions, geological composition, and the serpentinization process. The presence of alkali and alkaline earth metals in all sites (Table 1) indicates their vital roles in cellular processes, enzymatic reactions, and structural functions(Midgley, 1975). All sites also exhibited the presence of key transition metals, including manganese (Mn), iron (Fe), nickel (Ni), and copper (Cu). These metals play a crucial role as cofactors for enzymes that facilitate essential cellular activities, including

**Table 1:** Elemental analysis of water samples from different sites using inductively coupled plasma mass spectrometry (ICP-MS)

| Elements | Different Sampling Sites |  |  |  |  |
| --- | --- | --- | --- | --- | --- |
|  | Site 1<br>(µg/L) | Site 2<br>(µg/L) | Site 3<br>(µg/L) | Site 4<br>(µg/L) | Site 5<br>(µg/L) |
| Alkali Metals |  |  |  |  |  |
| Na | 110667.66 | 110667.66 | ND | ND | 82020.03 |
| K | 1150.4105 | 1150.4105 | 1299.455 | 5122.07 | 1356.67 |
| Rb | 0.096 | 0.096 | 0.127 | 0.28415 | 0.0155085 |
| Cs | ND | ND | ND | 0.0368 | 0.045 |
| Alkaline Metals |  |  |  |  |  |
| Mg | 444.32 | 444.32 | 33103.27 | 775.235 | 45443.144 |
| Ca | 25941.86 | 25941.86 | 20627.62 | 710.855 | 5329.415 |
| Sr | 3.055 | 3.055 | 2.4775 | ND | 2.381 |
| Ba | 0.945 | 0.945 | 0.5261 | ND | 5.04115 |
| Transition Metals |  |  |  |  |  |
| V | ND | ND | 0.1555 | 2.755 | 0.1616 |
| Cr | 0.198 | 0.198 | 1.47 | ND | 1.3545 |
| Mn | 7.19 | 7.19 | 12.95 | 0.647 | 18.2275 |
| Fe | 21.805 | 21.805 | 53.495 | 39.405 | 85.728 |
| Co | ND | ND | ND | ND | 0.1915 |
| Ni | 0.38 | 0.38 | 0.3415 | ND | 11.7385 |
| Cu | 0.9385 | 0.9385 | 0.8455 | 2.198 | 0.53525 |

Furthermore, variability in the concentrations of Total Organic Carbon (TOC), Total Carbon (TC), Non-Purgeable Organic Carbon (NPOC), and Total Nitrogen (TN) across the sampling sites (Table 2) has been observed. These findings provide valuable insights into the dynamics of organic matter, carbon cycling, and nutrient availability within the aquatic ecosystem. Variations in NPOC, TC, and TN across the sites may reflect differences in biological activity, the input of organic matter, and nutrient processing within the sampled environments.

**Table 2:** Physicochemical parameters of different water sampling sites.

| Parameters | Different Sample Sites |  |  |  |  |
| --- | --- | --- | --- | --- | --- |
|  | Site 3 | Site 4 | Site 1 | Site 2 | Site 5 |
| Temperature (°C) | 15.5 | 14.8 | 15 | 15 | 15.2 |
| pH | 7.48 | 10.72 | 7.98 | 7.6 | 7.59 |
| Methane (µM) | ~50 | ~75 | ~20 | ~15 | ~15 |
| Hydrogen ( µM) | ~ 173 | ND | ~153 | ~126 | ~235 |
| NPOC (µg/L) | ~38 | ~10 | ~6 | ~7 | ~8 |
| Total Carbon (TC) (mg/L) | ~90 | ~75 | ~50 | ~55 | ~75 |
| Total Nitrogen (TN) (mg/L) | ~0.6 | ~0.75 | ~0.2 | ~0.3 | ~0.5 |
| Nitrate (mg/L) | 0.314 | 0.59 | 0.123 | 0.102 | 0.407 |
| Sulphate (mg/L) | 2.39 | 27.75 | 3.1 | ND | ND |
| E<sub>h</sub> (mV) | -143 | -225 | -183 | -167 | -98 |

### 3.2 Morphological pattern of the microbial population from different sites-

The scanning electron microscopic images provide significant information about the morphological patterns of the microbial populations at the sampling sites (Figure 1). The highest microbial population was observed in the Site 1 sample. Most of the microbes were rod-shaped and varied in size. Additionally, another round, oval-shaped microbial population was found, highlighting the microbial diversity of microbes. The site 4 shows a considerably lower abundance of microbes, while the site 3 has a lower microbial population than the site 4. Individual scattered rod-shaped microbes of varying sizes are present at both site 3 and site 4, along with a clustered cell population. The site 5 demonstrates the presence of filamentous fungal or algal cells, as well as other bacteria.

### 3.3 Community divergence pattern across different sites based on Shotgun Metagenomic Sequencing analysis

Community divergence pattern of different sites were analysed based on the alpha and beta diversity metrics to compare the microbial community symmetry, richness, and composition as well (Figure 2). The site 5 shows the highest microbial diversity, as indicated by the diverse colors at the species level among all five sites (Fig. 3S). The Bray-Curtis dissimilarity matrix for beta diversity analysis was used to assess community composition between different sites (Figure 2a). The 3-dimensional principal coordinate analysis (PCoA) plot was created using the matrix of pairwise distances between samples calculated by the Bray-Curtis dissimilarity at the genus level (Figure 2b). Water samples with similar microbial composition profiles are closer to each other, while samples with different profiles are farther away.

**Figure 3.**
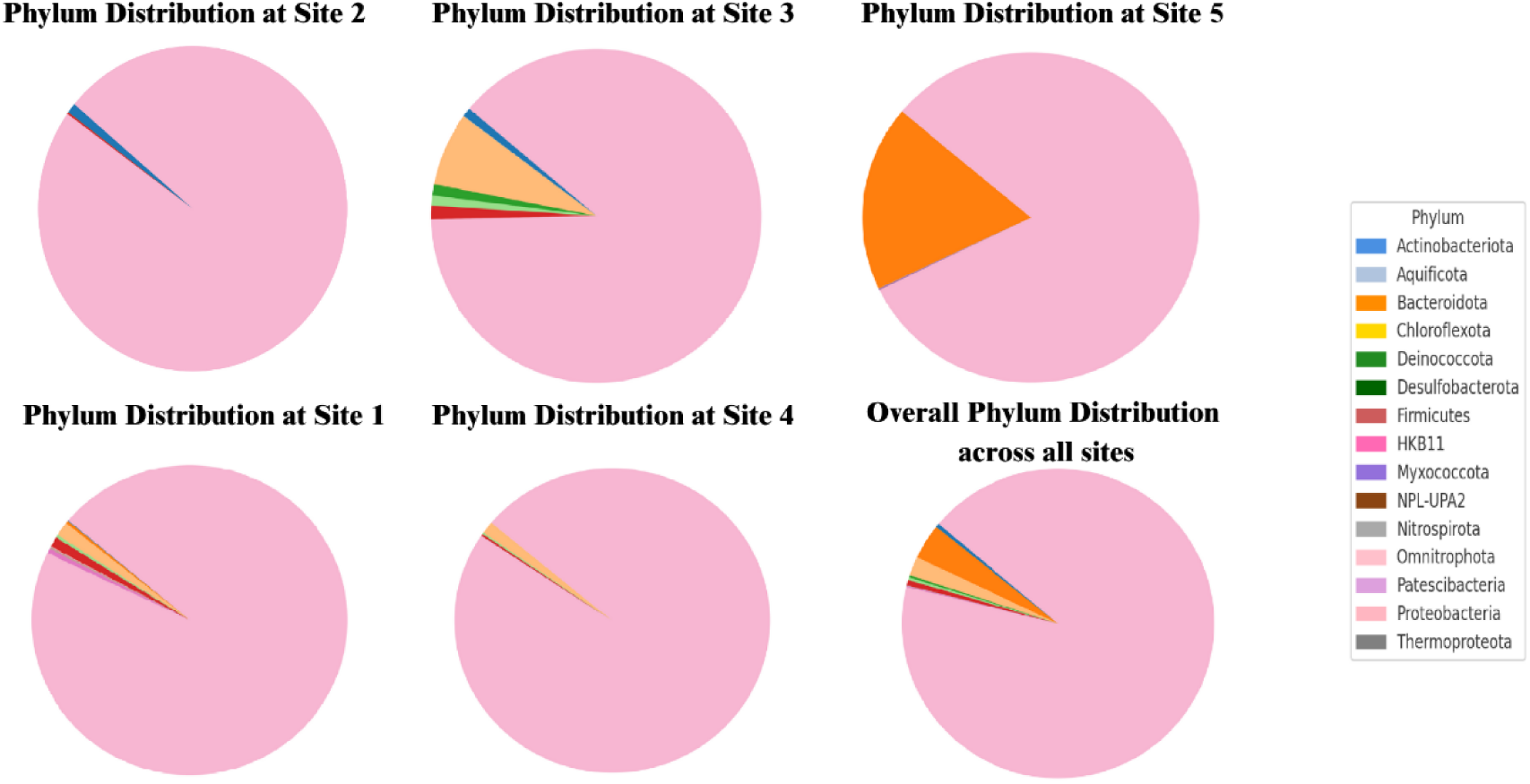
Phylum-level distribution pattern across sampling sites

Furthermore, considering the different phylum-level distribution pattern across the different sites, the highest overall abundance pattern was found for the taxonomic profile phylum Proteobacteria with a coverage of 92.7% (Figure 3). It had highest relative abundance of 98.8% in Site 2, with a lowest relative abundance of 81.7% atsite 5. It includes a varied distribution of betaproteobacteria, deltaproteobacteria and gamma-proteobacterial population across the different sites. The next most abundant phylum was found to be Bacteroidota with an entire abundance of about 3.7%. It is comprised with the relative abundance of 0.3 and 18.1 % of in the Site 1 and site 5 respectively. Then the third highest comes to the phylum Chloroflexota and distribution in Site 4, Site 1 and site 5 is 1.3%,1.5% and 7.2% respectively. Desulfobacterota counts the total abundance of 0.3% with a relative distribution of abundance is 0.2%, 0.3%, and 1% in Site 4, Site 1and Site 3, respectively. Firmicutes covers a total abundance of 0.2% with a relative abundance pattern of 0.6%, 0.4%, 0.2% in Site 1, Site 3 and Site 2, respectively. The maximum abundance of Euryarchaeota was found in Site 2, whereas Crenarchaeota was only in site 5. Additionally, the in-depth metagenome analysis reveal the presence of methanogenic archaea not reported before from the water sampling sites (Table 1S Excel file).

### 3.4 Microbial correlation with environmental variables –

The Non-metric Multidimensional Scaling (NMDS) plot illustrates the ordination of the listed microbial population structure based on Bray-Curtis dissimilarity, along with aligned phylum centroids and environmental vectors (Figure 4). The microbial population group is marked throughout the sites (indicated by a black star). Here, site 5, site 1 and site 2 hold a more discrete position in the ordination area, displaying varying microbial compositions compared to the strictly grouped sites of site 3, site 4 and site 2. This arrangement supports the influence of environmental factors on microbial populations at these sites. Red arrows denote environmental slopes that are significantly associated with microbial population diversity. The elongated vectors, such as Mn, Fe, Ba, Na, and total carbon (TC), show the maximum effect on microbial populations. The figure shows Methane, copper (Cu), chromium (Cr), and calcium (Ca) have a considerable, although reasonable, effect on microbial dispersal. Colored circles denoting phylum centroids are mapped to show the regular NMDS locations of individual phylum incidences across sites. Firmicutes, Proteobacteria, Chloroflexota, and Actinobacteriota are centrally positioned, indicating extensive spreading through the sites. Deinococcota and Aquificota are positioned at more peripheral sites, possibly exhibiting niche particularity toward individual gradients. The even labelling and colouring confirm that each phylum is denoted only once, avoiding overemphasis on profuse taxa such as proteobacteria. The cumulative R² value from the environmental fitting is 8.95, signifying a strong effect of environmental factors on microbial communities. The discrete environmental vectors displaying substantial r² data confirm the robustness of the data fitting. The potent effects of elements like Mn, Fe, and Na, as well as methane concentration, emphasize the roles of both redox-associated and geochemical factors in influencing microbial bionetworks in these sites. The analysis shows that the community structure is established with a substantial abundance of nickel (Ni) and iron (Fe), supported by the length and direction of individual vectors. Nickel and iron act mutually at the core of hydrogen metabolism; nickel acts as a catalytic centre, and iron acts as an organisational stabilizer with an electron transporter, creating a functional pair that is equally early and indispensable in biotic workflow. In the ordination, the presence of the iron (Fe) vector is most distinct, showing an effective association value (r² = 0.432) with variations in microbial populations. The iron vector extends through the ordination area toward taxa displaying Nitrospirota, Proteobacteria, and Firmicutes, implying an effective association among these taxa with the presence of iron at the different sites. This pattern aligns with the metabolic potency of these taxa, particularly highlighting their involvement in hydrogen-metabolizing enzymes. Locations such as site 2 and site 4, situated in close proximity to the iron vector, suggest that iron influences these sites comparatively. A significant presence of Ni, characterized by its sharp vector length and correlation value (r² = 0.401), also supports the effectiveness of nickel (Ni) containing hydrogen and methane-metabolizing enzymes (Prakash et al., 2014; Tianwei Wang1, 2019). The maximum abundance of Na, Sr, and Cu was found at the site 2. The study revealed that in methanotrophic bacteria, copper is involved in the functioning of methane monooxygenases, which are essential for methane oxidation (Tucci and Rosenzweig, 2024). A high abundance of the phylum Proteobacteria at the site 2 supports the presence of methanotrophs and the utilization of copper from the environment (Kenney and Rosenzweig, 2018). Proteobacteria, comprising various halotolerant microbes, are well-suited to the site 2 due to the abundance of sodium. Certain halotolerant sulfate-reducing bacterial populations under the Desulfobacteriota phylum can also be supported by high sodium content (Thompson et al., 2022). Members of the phylum Firmicutes have also been isolated from saline environments(Seong et al., 2018). A higher NPOC supports the heterotrophic microbial growth of the phylum Firmicutes, while TC favors the presence of both autotrophic and heterotrophic metabolism in some proteobacteria (Sui et al., 2024; Zheng et al., 2022). A lower TN restricts the nitrification and denitrification processes by reducing the respective microbial populations (Chen et al., 2022). Considering the abundance of Firmicutes in site 1, a study by Fuhren et al., 2021 found a strong positive growth impact of calcium on the microbial community of Firmicutes, correlating strongly with calcium abundance at thesite 1 (Fuhren et al., 2021). The study found that the presence of nitrate and sulfate favors the growth of proteobacteria, which are abundant at thesite 1 (Li et al., 2021). Research shows the abundance of microbial lineages under the phylum Desulfobacteriota and Chloroflexi in high sulfate ion concentrations (Elshahed et al., 2003; Magnuson et al., 2023). A comparatively high vanadium concentration can be tolerated by the microbial communities of Proteobacteria and Firmicutes, which supports their abundance pattern at the site 4 (Zhang et al., 2019). The metagenomic study on the role of Chloroflexi proved their adaptability to various carbon-rich sources (Narsing Rao et al., 2022). Desulfobacteriota was found to be present at the site 4 rich in carbon and nitrogen sources necessary for their active metabolism (Zhao et al., 2024). Furthermore, the presence of methane across the sampling sites significantly correlates with the presence of methanogenic archaea observed through in-depth sequencing.

**Figure 4.**
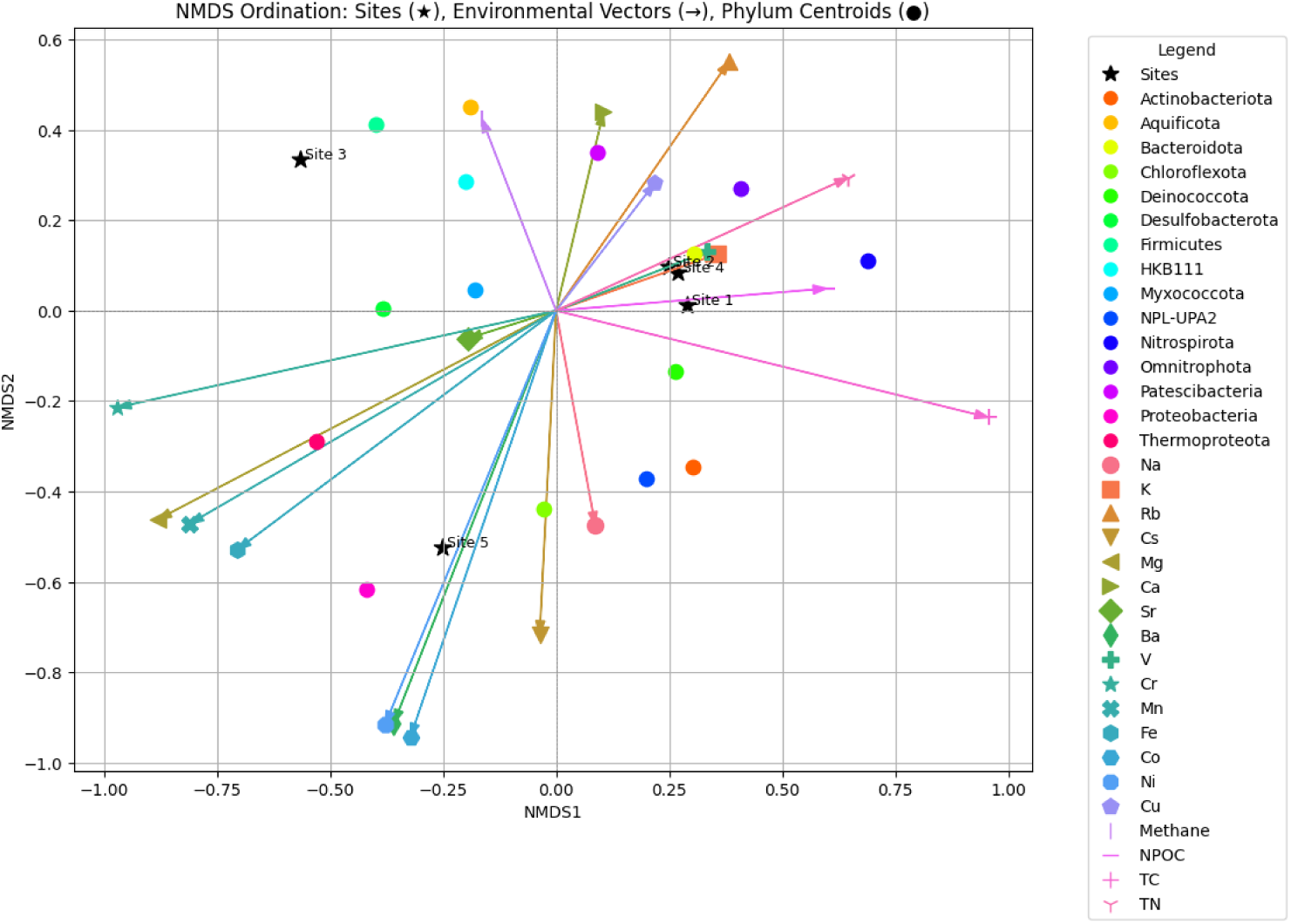
NMDS analysis of microbial community composition across sampling sites

### 3.5 Taxonomic comparative view of the most abundant phylotype across sites and microbial co-occurrence

An operational taxonomic unit (OTU) based study of sampling sites gives us more details on the distribution pattern based on genetic similarity (Figure 4S). The taxonomic abundance heatmaps with sample clustering show the highest abundance of diverse microbes at different sites (Figure 4S and supplementary information).

The cooccurrence study shows a positive correlation profile between the combination of two taxa, having a correlation value of 1 (Table 2S and Figure 6), which signifies an absolute positive linear relationship between two taxa of these sites. Results show that the presence of a combination of taxon such as CSP1-1-Bacteroides, Schlegelella_A - PNNF01, Rubrivivax-Giesbergeria, Rubrivivax-Kinneretia, Rubrivivax-Pelomonas, Rubrivivax-Sphaerotilus, Rubrivivax-Vogesella, Rubrivivax-Methyloversatilis, Rubrivivax-Brevundimonas, Rubrivivax-Elstera is highly favourable by the means of statistical analysis. The zero-*p* value indicates the likelihood of co-occurrence by chance is very little, reinforcing the potency of the spotted relationship. These co-occurrences suggest the probable syntropy or shared environmental fondness. In contrast, a negative correlation between taxa implies their probable antagonistic relationship or niche-related struggle. The combination of these taxa such as Cutibacterium-Stenotrophomonas, Cutibacterium-Meiothermus_B, Cutibacterium-Escherichia, Acidovorax-Aeromonas, Aeromonas-Flavobacterium, Aeromonas-Macellibacteroides are found to be negatively correlated with each other with the support of statistical analysis (P <1). In the heat map showing microbial co-occurrence across the sites, the maximum co-occurrence is represented in red, whereas lower one is indicated in blue.

**Figure 6.**
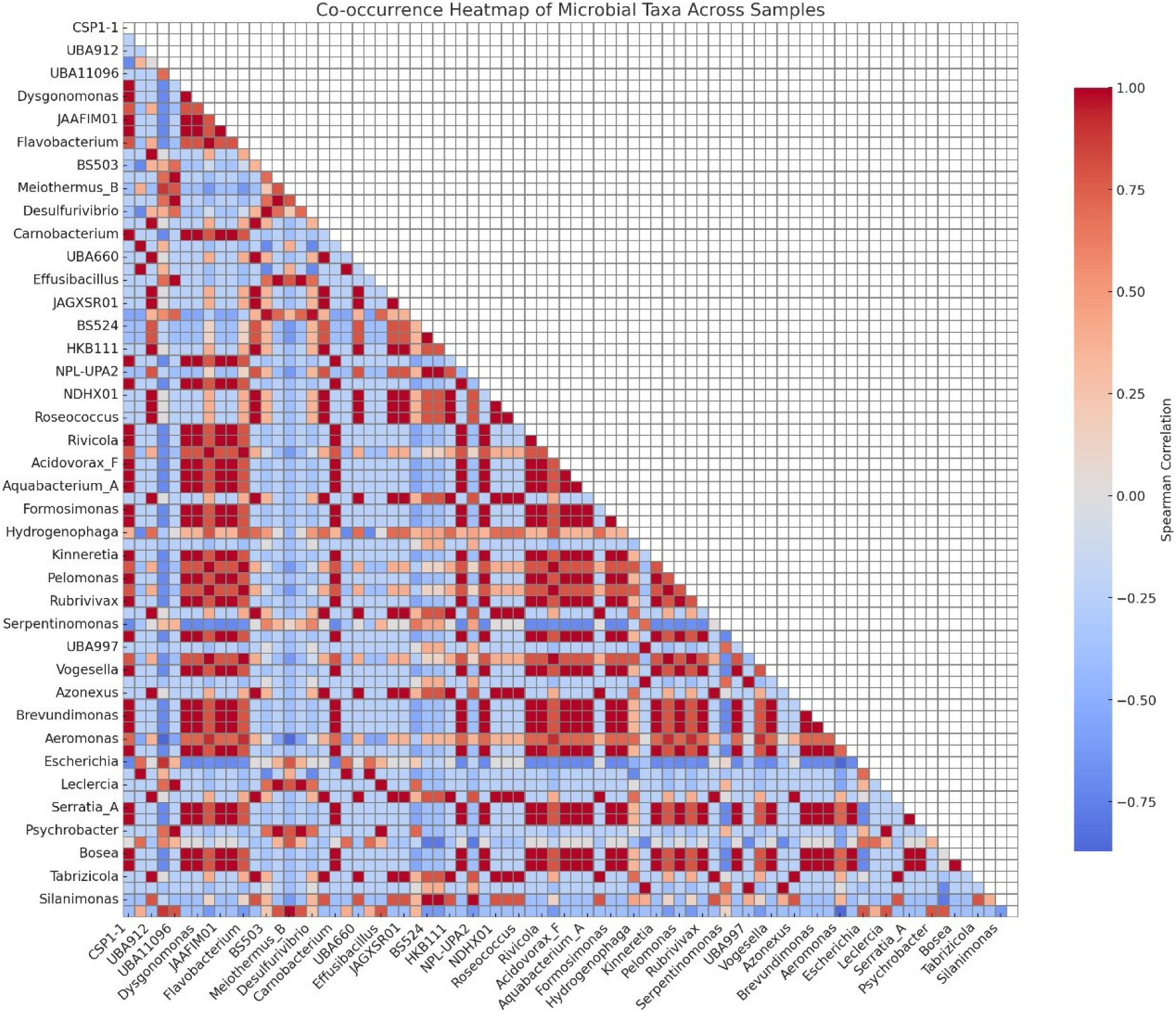
Microbial co-occurrence profile across the water sampling sites.

### 3.6 Microbial Consortia

The metagenomic study reveals a diverse microbial consortium across different sites, comprising hydrogen producers, utilizers, and methanogens. Each of them contributes towards making the biogeochemical pathway functional. The abundance of chloroflexi and methanogens in different water sampling sites suggests a possible syntrophic relationship that enables them to collaboratively degrade organic matter, resulting in methane production as the end product, as reported previously (Bovio-Winkler et al., 2023). Furthermore, the presence of Dehalococcoidia bacteria that can utilize environmental hydrogen for reductive dehalogenation and may aid in degrading anaerobic hydrocarbons in sulfate-reducing environments at elevated temperatures at the water sampling sites. This indicates their significant role in the anaerobic oxidation of specific compounds (LU, 2006. ; Zehnle et al., 2023). Interestingly, the metagenomic study of the serpentine-hosted tableland and lost city has not found this group of microbes in their reporting (Brazelton et al., 2012). Furthermore, *Thermorudis* sp. (0.1%) can oxidize carbon monoxide, playing a significant role in the geochemical cycle by contributing substantially to the nitrogen cycle through denitrification (King and King, 2014; Mefferd et al., 2022). *Thermus* sp. can coexist with thermophilic archaea, forming a microbial association that leads to complex interrelationships within an environmental niche (Coman et al., 2013).

Desulfurivibrio is an essential participant in the nitrogen and sulfur cycles at site 3 and site 4. The high concentration of sulfate ions, acting as an electron acceptor (Table 1), enhances the growth and metabolic activity of Desulfurivibrio, influencing the transformation and utilization of sulfur compounds at the site 3 and site 4 (Li et al., 2022). *Hydrogenophaga* sp., known as mixotrophs, utilize hydrogen for autotrophic growth and make up 0.9% of the population at the site 3; they are a commonly found genus in serpentine sites (Suzuki et al., 2013). *Psychrobacter* sp., present at 0.1%, is adapted to cold environments and plays a role in the carbon cycle (Bakermans, 2018), while *Stenotrophomonas* sp., also at 0.1%, contributes to the sulfur and nitrogen cycles with sulfate and nitrate as terminal electron acceptors (Chauviat et al., 2023), (Mukherjee and Roy, 2016). The site 4 exhibits a similar pattern of microbial population, differing only in their identified counts. Notably, *Tepidomonas* sp. was identified here, supporting the growth of the Chloroflexi group by producing indoleacetic acid and pantothenic acid (Xian et al., 2020).

### 3.6 Metabolic Pathway Abundance and gene family

The metabolic pathway abundance heat map with sample clustering shows 75 most abundance patterns of functional pathways as per the colour gradient such as glycolysis, pyruvate fermentation, fatty acid metabolism, different amino acids and nucleotides synthesis (Fig. 4S).

The heat map of the gene family of different sites clustering shows gene abundance patterns as per the colour gradient (Figure 5S). Rubredoxin (UniRef90_A0A060NPR2) that is involved in the iron clustering and electron transport was found to be most abundant in the site 4 and then the site 3, related to the maximum abundance of *Serpentinomonas raichei* (Lu and Imlay, 2021). The protein is an electron carrier in the anaerobic environment that supports anaerobic metabolism of *Serpentinomonas* sp. as per its distribution across the serpentinite sites. Rubredoxin and its redox shuttling property provide a molecular analogue to the redox practice innate to serpentine surroundings. Both approaches depend on precise electron transport between redox associates. Rubredoxin could be imperative for the microbes harboured in serpentinization environments, allowing them to tap the profuse hydrogen and exploit the Fe2+/Fe3+conversion for metabolic courses influenced by oxidation-reduction of this bionetwork. Cbb3-type cytochrome oxidase, subunit 3 (UniRef90_A0A060NP58) that corresponds to the source of *Serpentinomonas raichei* is prevalent in the maximum part in site 3 than in site 4. The Cbb₃-type cytochrome oxidase enables microaerophilic growth of *Serpentinomonas raichei* by maintaining low cytoplasmic O₂ tension, thereby protecting and activating its O₂-sensitive enzymes (Suzuki et al., 2014; Weingarten et al., 2008; Zhang et al., 2023). The maximum presence of hemin uptake protein (UniRef90_A0A060NRG4) in site 4 supports the sequestering of iron from the environment, supporting the maintenance of Ni, Fe-hydrogenase (Group2b) in *Serpentinomonas raichei* (Suzuki et al., 2014). In Clostridium difficile, 2-oxoisovalerate dehydrogenase (UniRef90_A0A060NH86) balances redox equivalents and drives ATP synthesis under anaerobic conditions; by analogy, it may similarly promote autotrophic growth in *Serpentinomonas raichei* (Kim et al., 2006). Consequently, the prevalence of 2-oxoisovalerate dehydrogenase in thesite 3 and site 4 supports this finding.

The presence of hydrogenic *Serpentinomonas* sp. across the sites supports the existence of a hydrogenase gene required for hydrogen metabolism. This observation is substantiated by the RefSeq sequence of *Serpentinomonas* sp. 000696225, which indicates the presence of coenzyme F420-reducing hydrogenase, alongside the Ni, Fe-hydrogenase enzyme, maturation factors, and formate dehydrogenase, thereby reinforcing its ability for hydrogen utilization, comparable to other Serpentinomonas species, as previously elucidated (Hendrickson and Leigh, 2008).

A very low abundant NPL-UPA2 bacterium Unc8 shows a close identity with GCA_003574845.1 of more than 99%, including genome coverage of 33.89% (abundance hit 0.0002) and 24.41% (abundance hit 0.00037) in site 4 and site 1, respectively. The genome assembly reveals the presence of different subunits of CO dehydrogenase/acetyl-CoA synthase complex (cdh), the formate dehydrogenase (Fdh), sulfurtransferase, and the bifunctional methylenetetrahydrofolate dehydrogenase/methenyltetrahydrofolate cyclohydrolase FolD (folD) (Suzuki et al., 2018). These enzymes are all involved in the Wood-Ljungdahl Pathway, which aids in acetate formation by converting carbon monoxide and coenzyme A into a methyl group, thus enhancing the metabolic functions of the acetogenic NPL-UPA2 bacterium Unc8. *Desulfurivibrio* sp004332195 shows a close identity with GCA_004332195.1 of more than 98%, including varied genome coverage in site 4, site 1,site 5 and site 3. The genome assembly encodes dissimilatory sulfite reductase subunits α (dsrA) and β (dsrB), confirming the organism’s capacity for sulfate reduction.

Furthermore, the prevalence of KEGG-annotated genes associated with the most widely distributed methanogenic archaea, *Methanosarcina acetivorans C2A*, observed across all sites illustrated in Figure 7, reveals genes that are involved in essential metabolic pathways, particularly those pertinent to methane metabolism. These genes encompass heterodisulfide reductase subunits, succinate dehydrogenase, and tetrahydromethanopterin S-methyltransferase, each exhibiting variable abundance patterns across distinct samples. This plot showcases the 25 most abundant genes of *M. acetivorans C2A*, selected based on their highest average RPKM across all samples.

**Figure 7.**
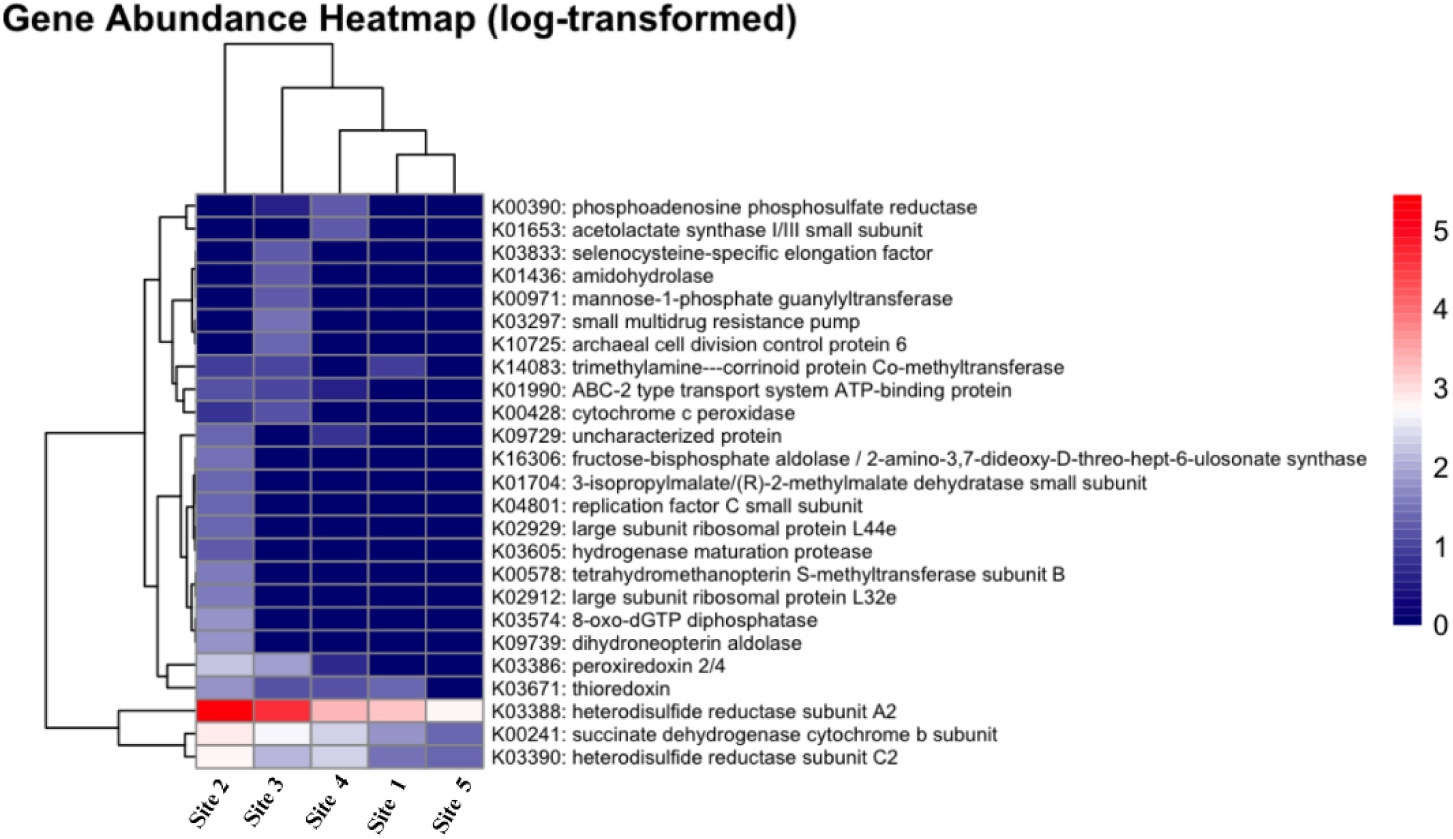
The heat map of the gene family shows gene abundance patterns of *Methanosarcina acetivorans* C2A as per the colour gradient.

### 3.7 Diversity of Hydrogenase

Hydrogenases are metalloenzymes that favour the reversible molecular hydrogen oxidation. Based on metal cofactors and conserved motifs, they are typically categorized into different classes like [Ni-Fe], [Fe-Fe] and [Fe] hydrogenase (Søndergaard et al., 2016). Here, we study the phylogenetic association among the sequences similar to hydrogenase without allocating them to traditional group categorization, giving an unbiased sight of their evolutionary variance. Phylogenetic analysis of the hydrogenase gene within the microbial community indicates the likely presence of hydrogenase-bearing sequences and highlights microbial divergence concerning the [Ni-Fe] hydrogenase sequence across the different sites. This study covers site 1 and site 2 together, site 3 and site 4 together site 5. The hydrogenase enzyme-based phylogenetic tree presents several well-verified clades on the basis of sequence similarity.

The phylogenetic tree of the site 1 and site 2 showed (Figure 8) the presence of various microbial species within the ancestral clade of the Proteobacteria Group 1, along with five most recent common ancestors (MRCA). Species of *Syntrophus* and *Shewanella* come as sister taxa with differences in their Ni-Fe hydrogenase sequence under MRCA. *Escherichia* sp, being a closest relative to *Syntrophus* and *Shewanella*, under the common node of gammaproteobacteria, shows significant divergence of Ni-Fe hydrogenase. Species under the Betaproteobacteria and Betaproteobacteria 2 show differences in their hydrogenase. Species under the Proteobacteria group 1, Betaproteobacteria2, Nitrosomonadles, Deltaproteobacteria, being a member of the common node, share a common sequence with significant divergence as per the distance from each other. *Clostridia* sp having Fe-Fe hydrogenase (involved in anaerobic hydrogen production) set as an outgroup participant to the phylogenetic tree with a promising divergence from the Ni-Fe hydrogenase sequence (Calusinska et al., 2010).

**Figure 8.**
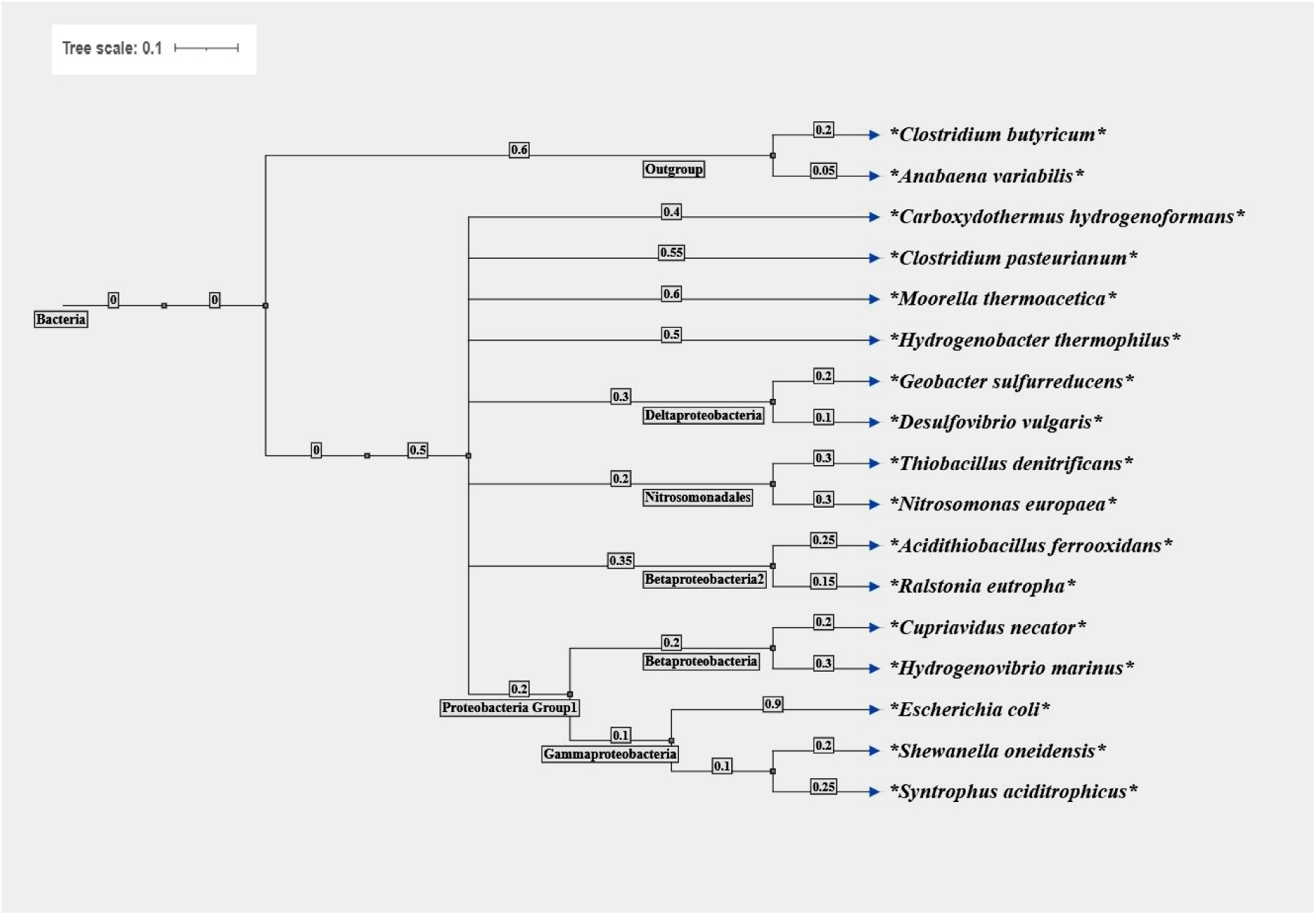
Phylogenetic diversity of putative [Ni-Fe]-hydrogenase spotted in metagenomes from site 1 and site 2

In the case of the site 3 and site 4, although the anaerobic and proteobacteria Group 1 share a common monophyletic group, they possess different species with significant divergence among them (Figure 9). The metagenome assembled genome (MAG) of UBA 997 sp002291945, identified from an environmental sample carries a hydrogenase enzyme that is distinctly related to hydrogen utilizing Ni-Fe hydrogenase gene of *Serpentinomonas* sp, *Hydrogenophaga* sp. This genomic variety is identified as an uncultured bacterial and archaeal species with limited phenotypic data compared to genomic data. Species under MRCA come as sister taxa share maximum sequence similarity and this helps us to get the closeness of *Serpentinomonas* sp and *Hydrogenophaga* sp based on hydrogen utilization. GPS1B09 sp18725345, JAGXSE01 sp018333865, and JAGXSE01 sp018333405 are the MAG genomes identified from environmental samples, known to possess Ni-Fe hydrogenase gene that are significantly different from those *Serpentinomonas* sp and *Hydrogenophaga* sp. GPS1B09 sp18725345, JAGXSE01 sp018333865 are the microbial culture having limited phenotypic data than genomic one. The Anaerobic divergence of the proteobacteria clade, having UBA997 and *Desulfurivibrio* sp. shares the most common Ni-Fe hydrogenase sequence among others in this phylogenetic tree.

**Figure 9.**
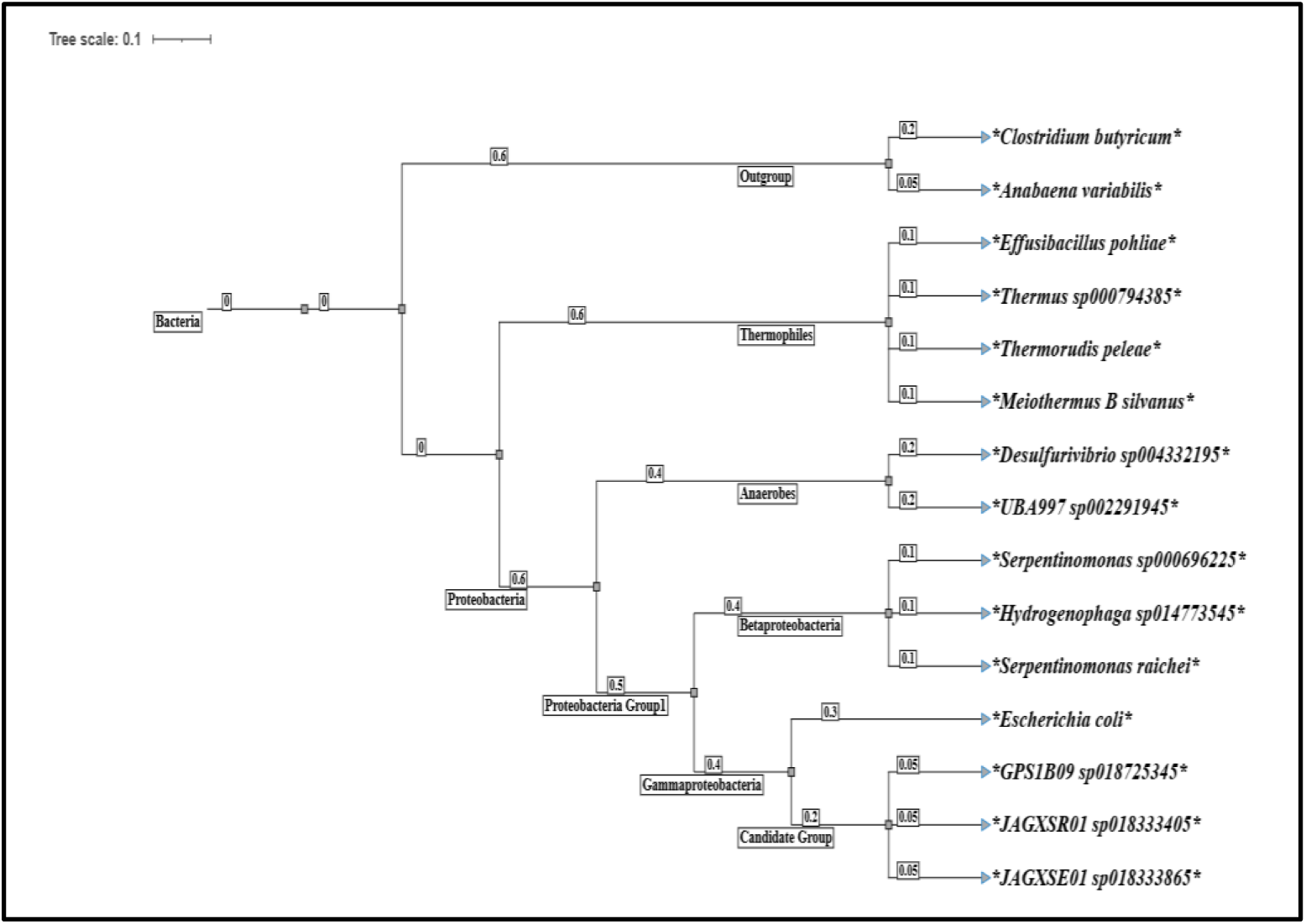
Phylogenetic diversity of putative [Ni-Fe]-hydrogenase spotted in metagenomes from site 3 and site 4.

In site 5 two of the clades proteobacteria and proteobacteria group 1, hold all of the species with four MRCA (Figure 10). The sister taxa under beta and gammaproteobacteria show maximum closeness among their Ni-Fe hydrogenase gene sequence, whereas they are considerably divergent from the delta and alphaproteobacterial taxa. *Azospirillum* sp shows difference in Ni-Fe hydrogenase gene sequence than the other sister taxa of the same MRCA. *Desulfovibrio* sp shows significant divergence from the rest of the other taxa of the same clade of site 5.

**Figure 10.**
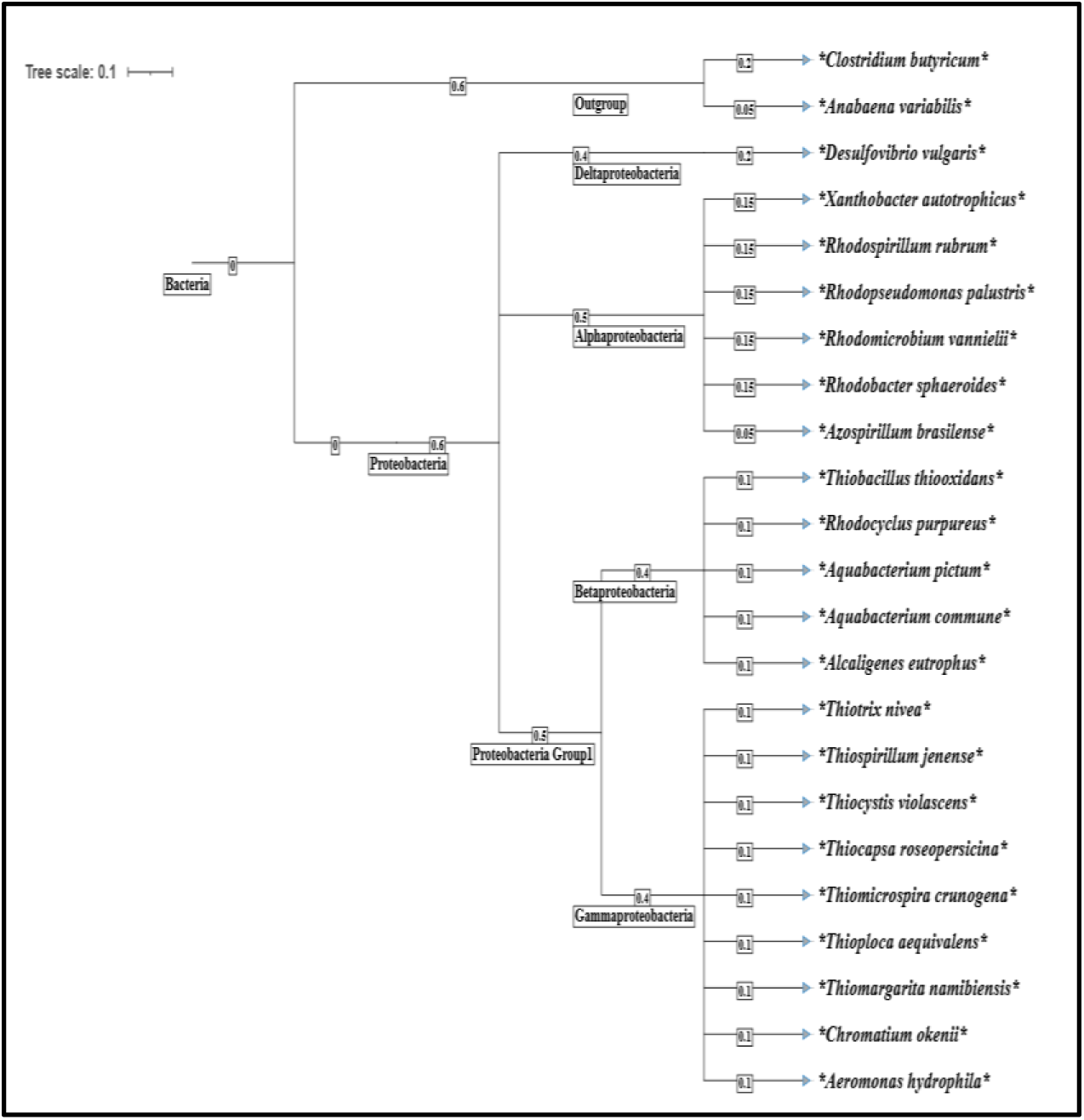
Phylogenetic diversity of putative [Ni-Fe]-hydrogenase spotted in metagenomes from site 5.

## 4. Summary and Conclusion

Serpentinization mafic rock systems serve as terrestrial analogs for Earth’s primordial atmosphere. The water sampling site in northern California sustains diverse microbial communities adapted to these early-Earth-like conditions. A well-balanced microbial population comprising both hydrogenogenic and hydrogenotrophic microorganisms, exhibiting notably varied hydrogenase gene abundance, was identified. The presence of methanogenic archaea, with *Methanosarcina acetivorans* being the most abundant at site 1, enhances the study’s significance in relation to relevant geochemical findings. Analysis reveals similarities among the microbial populations of Site 3, GPS site 4, and Site 1, while the microbial community at Site 5 is notably more complex. Site 5 demonstrates its distinctiveness through the presence of microbial phyla such as Bacteroidota (with higher abundance than at Site 1), Myxococcota, and Nitrospirota, each contributing unique functional properties that characterize this site distinctly.

### Future Prospects

Metagenomic analysis of water samples holds significant promise for identifying the diversity and functional capabilities of microbial communities, especially those involved in hydrogen metabolism. It highlights the fundamental role microbes play in the biogeochemical cycle, with an emphasis on sustainable energy pathways. The future prospects include harnessing these microbial processes to improve biohydrogen production, along with a strong focus on inhibiting hydrogen-consuming microbes to enhance overall hydrogen yield. These advancements could support improvements in renewable energy and conservation biotechnology.

## Acknowledgments

This study is supported by the Advanced Research Projects Agency-Energy (ARPA-E) through grant Award # DE-AR0001894.

