## Supplemental for "Exploring the Microbial Geobiological Pattern Across the Serpentinization Sites through Metagenomic and Elemental Composition Analyses": Supplementary File.pdf.pdf

**Content**

**Results and discussion**

**Figure 1S** Water sampling setup used for collecting samples

**Figure 2S (a): -** Sampling sites are marked with a yellow circle (**b & c**): - Different water sampling sites where samples were collected during this study.

**Figure 3S.** Composition Barplots illustrating the microbial composition at the Species level

**Figure 4S**. The metabolic pathway abundance heatmap with sampling site Clustering.

**Figure 5S.** The heat map of the gene family with sampling site clustering shows gene abundance patterns as per the colour gradient.

**Figure 6S** Archaeal abundance profile across different sites. Series 1, 2,3,4,5 relate to site 4, site 1, site 5, site 3, and site 2, respectively.

**Table 1S** Presence of methanogenic archaea from the water sampling sites.

In the supplementary Excel file.

**Table 2S.** Microbial Co-occurrence Analysis

**Results and discussion:**

**Community divergence pattern across the sites-** The alpha diversity was measured with the Shannon index. The genus level distribution pattern based on the alpha diversity metrics shows a Shannon index of 0.23137 (site 4 ), 1.376137 ( Site 1 ), 1.963993 (site 5), 0.836364 (Site 3), 0.873711 (Site 2). It shows a maximum evenness within the site 5(Fig. 3S). The bar plot of alpha diversity shows the presence of a maximum 40 genera in site 5(Provided in the supplementary Excel sheet).

In case of beta diversity, the microbial populations of spring sites on the PC2 axis, such as site 3, site 4, and site 1, have a distinct difference along the PC2 axis from each other, primarily due to the second leading source of alteration in microbial composition, accounting for approximately 34.78% of the dissimilarity. The microbial population at the Site 2 is considerably dissimilar along PC1, which is the largest basis of alteration, accounting for approximately 57.62% of the dissimilarity. The microbial population of the site 5 exhibits changes along both PC1 and PC2, which represent the two primary bases of dissimilarity, accounting for approximately 92% of the overall variation. Consequently, the microbial population at Site 2 is the most variable, showcasing the greatest dissimilarity. In contrast, the microbial populations of Site 3, Site 4, and Site 1 show the least differences along the PC1 axis but differ significantly based on PC2. The microbial population of site 5 is the most complex, influenced by both the PC1 and PC2 axes of dissimilarity

**Taxonomic comparative view of the most abundant phylotype across sites-** The genus *Serpentinomonas* sp (63.78%) in Site 3 and comparatively lower abundance of genus *Hydrogenophaga, Silanimonas, Desulfovibrio*. The rest of the microbial count falls below 5%. The metagenomic study shows, the whole genome sequence identity of *Serpentinomonas raichei* , *Serpentinomonas* sp000696225 , *Hydrogenophaga sp014773545, Silanimonas* sp018334015, *Desulfurivibrio* sp004332195 of 99.2115% (GCF_000828895.1), 95.1828% ( GCF_000696225.1), 96.6347% (GCA_014773545.1), 91.4285% (GCA_018334015.1), 99.2141 % (GCA_004332195.1), respectively. A high pH of about 10.7 supports the growth of *Serpentinomonas* sp. a maximum abundance. The isolation of *Serpentinomonas raichei* from the highly alkaline spring site at the water sampling sites favors its growth at highly alkaline environment [1]. The study of alkaliphilic gammaproteobacterium*, Silanimonas lenta gen. nov., sp. nov.* shares the possibility for the presence of *Silanimonas* sp in the Site 4, as a highly alkaline area [2]. *Hydrogenophaga* sp can be grown in this alkaline area as it is found to be a less stringent mixotroph compatible with a wide range of environmental factors. Alkaliphilic indigenous *Hydrogenophaga* sp that is a bioremediator of petroleum hydrocarbons, can survive in alkaline sediment and this genus can also be able to grow in lab-used media at this high pH range [3-5]. The presence of *Desulfovibrio* sp. in an alkaline environment supports its presence in Site 4. A low dissolved oxygen level is supposed to favor the growth of *Serpentinomonas* sp due to its ability to survive in sub-atmospheric levels of oxygen[1]. Anaerobic growth of *Desulfovibrio* sp. is quite a favourable one in this low level of dissolved oxygen. Though the basic functionality of SRB was found mostly in the range of neutral and slightly acidic environment but the study of Kumar et al. 2020 show that SRB do prefer the alkaline pH like a neutral one. The *Desulfovibrio* sp and *Silanimonas* sp. belong to the class of deltaproteobacteria. Whereas *Hydrogenophaga* sp is in the class of betaproteobacteria. *Sphingomonas aquatilis* that is an alpha proteobacterium is present in very little amount is the temperature of this site also supports the moderate growth of these different class of bacterial populations that may not be the optimum but support fairly [1, 3, 6]. Subsequently, the Site 4 is enriched with the consortia of different classes of bacteria.

Site 3 is mostly rich with *Serpentinomonas* sp. having *Serpentinomonas raichei* , *Serpentinomonas* sp000696225, BS503 sp018725425 with an abundance of 54.78% and

27.31% respectively. These are found to have the whole genome identity of 99.0820% (GCF_000828895.1), 95.1476% (GCF_000696225.1), 96.9754% (GCA_018725455.1). The rest of the microbial count falls below 5%. A relatively moderate abundance of *Methanosarcina acetivorans* and lower *Methanobrevibacter ruminantium* count was found in site 3 and site 4 (Table XS) that further supported by detection of methane form both sites ( Table 1)

Site 1 is mostly enriched with *Serpentinomonas raichei* with an abundance of 30.12% along with other *Serpentinomonas* sp. having an abundance of 28.1%. Genus *Hydrogenophaga* and *Silanimonas* also occupied their over there with comparatively lower percentages than genus *Serpentinomonas.* The rest of the microbial count falls below 5%. *Desulfovibrio* was found to be present there but in very low count (0.3%). Though the microbial population abundance comes with some of the same genus as the site 3 yet it is not the scenario for the whole spring site. The whole genome sequence identity percentage also varies with *Serpentinomonas raichei* *, Serpentinomonas* sp000696225 , *Hydrogenophaga sp014773545, Silanimonas* sp018334015, *Silanimonas* sp018333835, *Silanimonas* sp003449515, *Desulfurivibrio* sp004332195 of about 97.2657 (GCF_000828895.1), 96.2912 ( GCF_000696225.1), 98.6210 (GCA_014773545.1), 89.8448 (GCA_018334015.1), 89.9689 (GCA_018333835.1), 89.4743 (GCA_003449515.1) respectively. Hence, it shows a different percent identity for the whole genome for a similar genus that may lead to different strains and even species for this spring site. site 2 is highly populated with genus *Escherichia*, *Klebsiella*, *Pseudomonas* and with very little abundance of *Serpentinomonas* sp000696225 , *Hydrogenophaga* sp. The whole genome identity percentage for *Escherichia coli* JM109, *Klebsiella michiganensis*, *Pseudomonas_E peli*, comes with 99.2823 (zymo22 *),* 99.1143%( GCF_002925905.1), 97.1409% (GCF_900099645.1). Site 2 shows the highest abundance of *Methanosarcina acetivorans* , an model acetrotphic methanognes along with *Methanobrevibacter ruminantium..*

Site 5 is there with the highest microbial abundance of *Aeromonas* sp.(28.16%), then it comes to gradually *Macellibacteroides* sp (10.93%), *Paucibacter* sp., *Rhodoferax* sp. The rest of the microbial count falls below 5%. The metagenomic study of the whole genome sequence identity finds the similarity of *Aeromonas salmonicida, Macellibacteroides* sp018054455*, Paucibacter_A* sp005503085*, Rhodoferax* sp002163715 of about 98.0440 % (GCF_001643305.1), 96.9406 % (GCA_018054455.1), 95.2113% (GCA_005503085.1), 92.6275% (GCF_002163715.1). This site harbours the lowest count of *Methanosarcina acetivorans.*

**Metabolic Pathway Abundance and gene family-** The presence of the super pathway ( Fig. ) of adenosylcobalamin salvage from cobinamide was found to be predominant in site 4, supporting its involvement in the metabolism of *Serpentinomonas raichei,* allowing them to proficiently get and produce crucial cofactors, cobalamin from the environment[7]. The biosynthesis pathway $preQ_0$ is a abundant across site 4, site 3, and site 1, supporting the growth of microaerophilic and anaerobic growth. The archaeon, *Thermococcus kodakarensis*, uses $preQ_0$ for the modification in tRNA. Moreover, some anaerobic bacteria have genes that code for enzymes like QueE, FolE, QueD and QueC that participate in the biosynthesis of $preQ_0$ [8, 9].

Tabrizicola sp013824055 exhibits a close identity with Rhodobacter sp., (GCA_013824055.1) of nearly 89%, with genome coverage of 25.13% (abundance hit 0.003). The genome assembly reveals the presence of formate dehydrogenase subunit alpha, hydrogenase accessory protein HypB (hypB), hydrogenase expression/formation protein HypE (hypE), HypC/HybG/HupF family hydrogenase formation chaperone (hypC), and hydrogenase, which are involved in hydrogen metabolism. The presence of these genes in closely related strains supports the hydrogen-metabolizing capability of Tabrizicola sp013824055. *Aeromonas veronii* exhibits a close identity with *Aeromonas veronii* bv. veronii strain CECT 4257 (GCF_000820225.1), showing almost 94.39% (site 1) and 96.41% (site 5 ) identity, including genome coverage of

9.85% (abundance hit 0.001) and 69.50% (abundance hit 0.034). The Refseq sequence shows the presence of formate dehydrogenase and hydrogenase enzyme subunits along with other accessory proteins that participate in hydrogen metabolism. *Serpentinomonas* sp000696225 and *Serpentinomonas raichei* shows a close identity with GCF_000696225.1 and GCF_000828895.1 of more than 90% and 94% respectively in four different spring sites like site 4, site 1, site 5, site 3 and site 2.

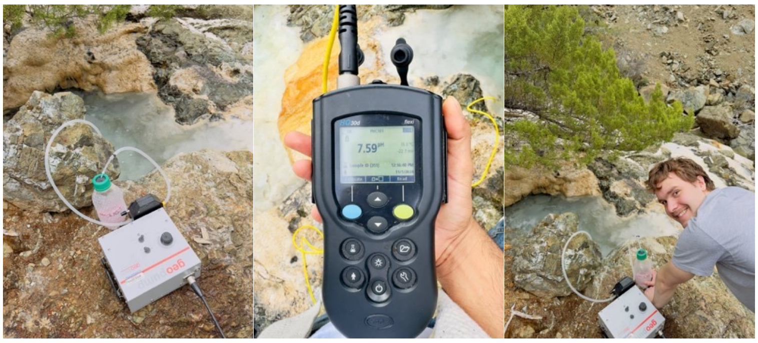

**Fig 1S.** Water sampling setup used for collecting samples

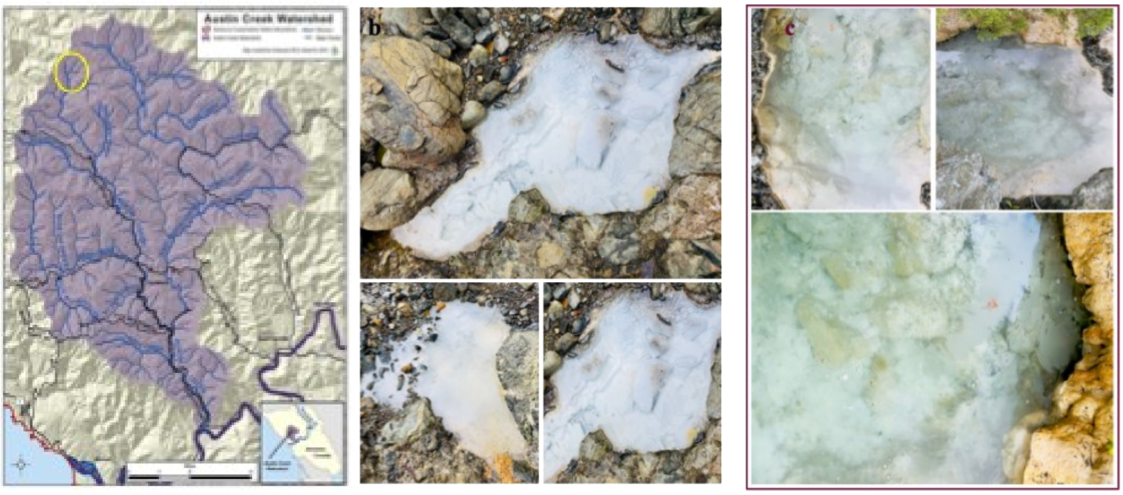


**Fig 2S (a): -** Sampling sites are with marked with a yellow circle (**b & c**): - Different water sampling sites where samples were collected during this study.

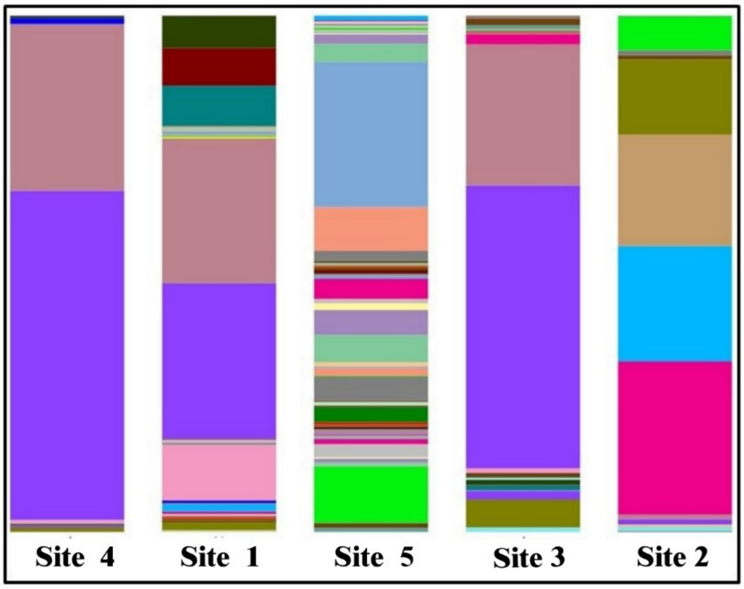


**Figure 3S.** Composition Barplots illustrating the microbial composition at the Species level

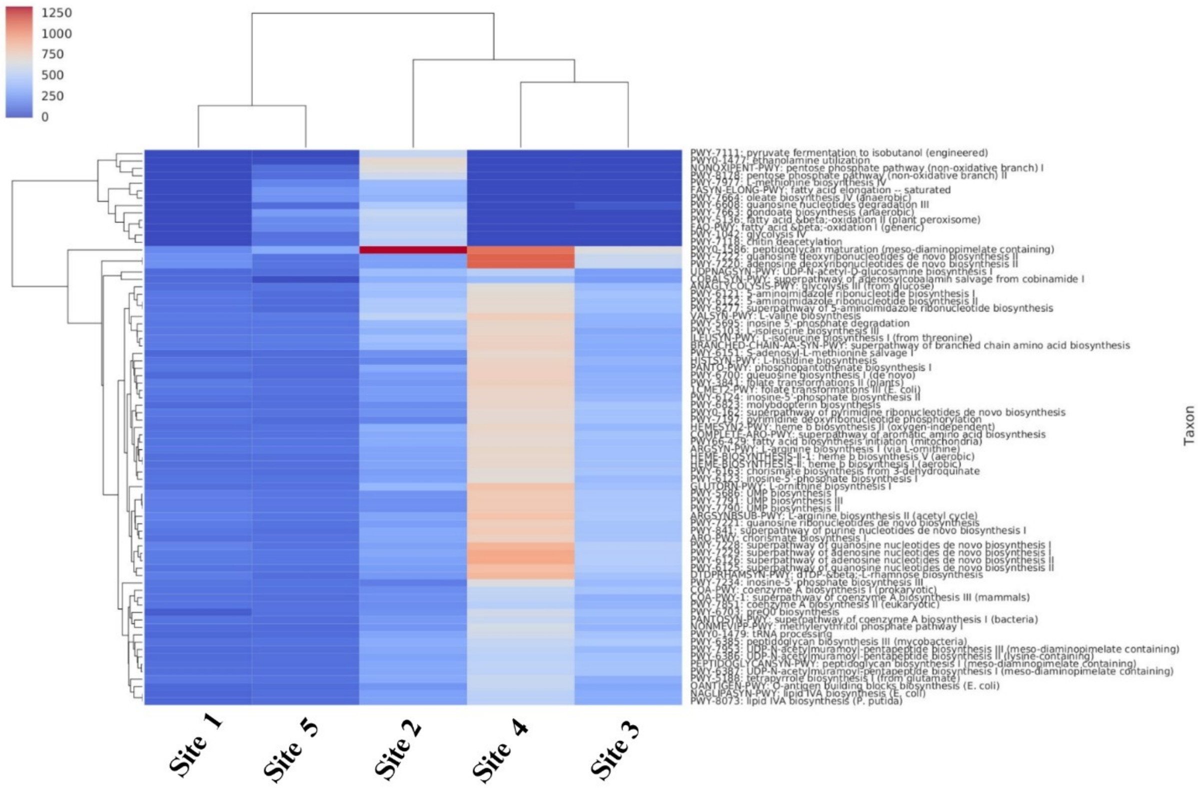


**Figure 4S**. The metabolic pathway abundance heatmap with sampling site Clustering. The higher abundance is represented in red, whereas the lower abundance is indicated in blue

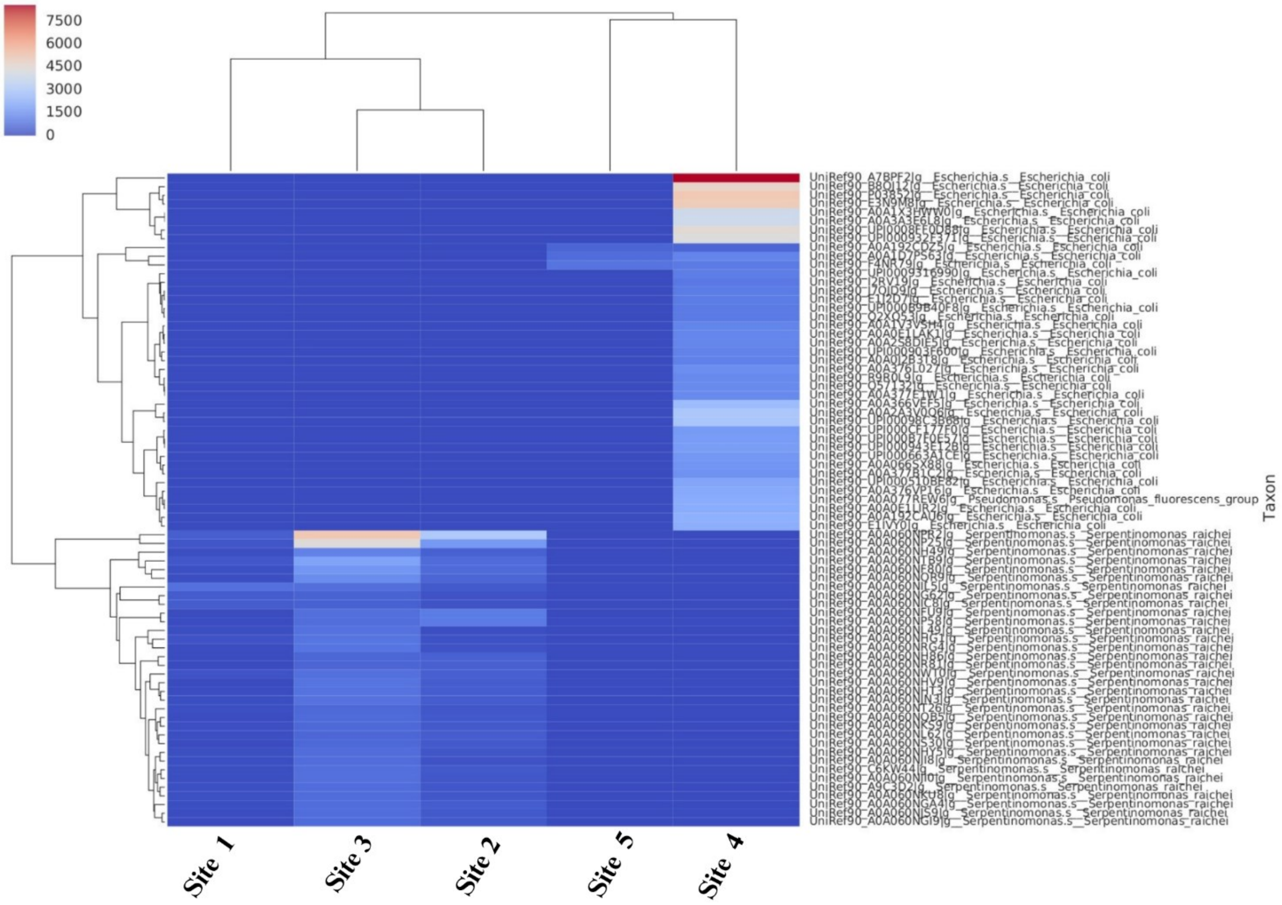


**Figure 5S.** The heat map of the gene family with sampling site clustering shows gene abundance patterns as per the colour gradient. The higher abundance is represented in red, whereas the lower abundance is indicated in blue.Each row represents the abundance for each gene family along with the species information, with the gene family ID shown on the right. Each column represents the abundance for each sample, with the sample ID shown at the bottom.

**Table 1S** Presence of methanogenic archaea  from the water sampling sites.
In the supplementary Excel file.

**Table 2S. Microbial Co-occurrence Analysis**

| Positive co-relation | | | |
|---|---|---|---|
| **Taxon 1** | **Taxon 2** | **Correlation** | **p-value** |
| **CSP1-1** | **Bacteroides** | **1** | **0** |
| **Schlegelella_A** | **PNNF01** | **1** | **0** |
| **Rubrivivax** | **Giesbergeria** | **1** | **0** |
| **Rubrivivax** | **Kinneretia** | **1** | **0** |
| **Rubrivivax** | **Pelomonas** | **1** | **0** |
| **Rubrivivax** | **Sphaerotilus** | **1** | **0** |
| **Rubrivivax** | **Vogesella** | **1** | **0** |
| **Rubrivivax** | **Methyloversatilis** | **1** | **0** |
| **Rubrivivax** | **Brevundimonas** | **1** | **0** |
| **Rubrivivax** | **Elstera** | **1** | **0** |

| Negatively Corelation | | | |
|---|---|---|---|
| **Taxon 1** | **Taxon 2** | **Correlation** | **p-value** |
| **Cutibacterium** | **Stenotrophomonas** | **0.89** | **0.0405** |
| **Cutibacterium** | **Meiothermus_B** | **0.89** | **0.0405** |
| **Cutibacterium** | **Escherichia** | **0.9** | **0.0374** |
| **Acidovorax** | **Aeromonas** | **0.92** | **0.028** |
| **Aeromonas** | **Flavobacterium** | **0.92** | **0.028** |
| **Aeromonas** | **Macellibacteroides** | **0.92** | **0.028** |

**Archaeal abundance pattern across the sites**

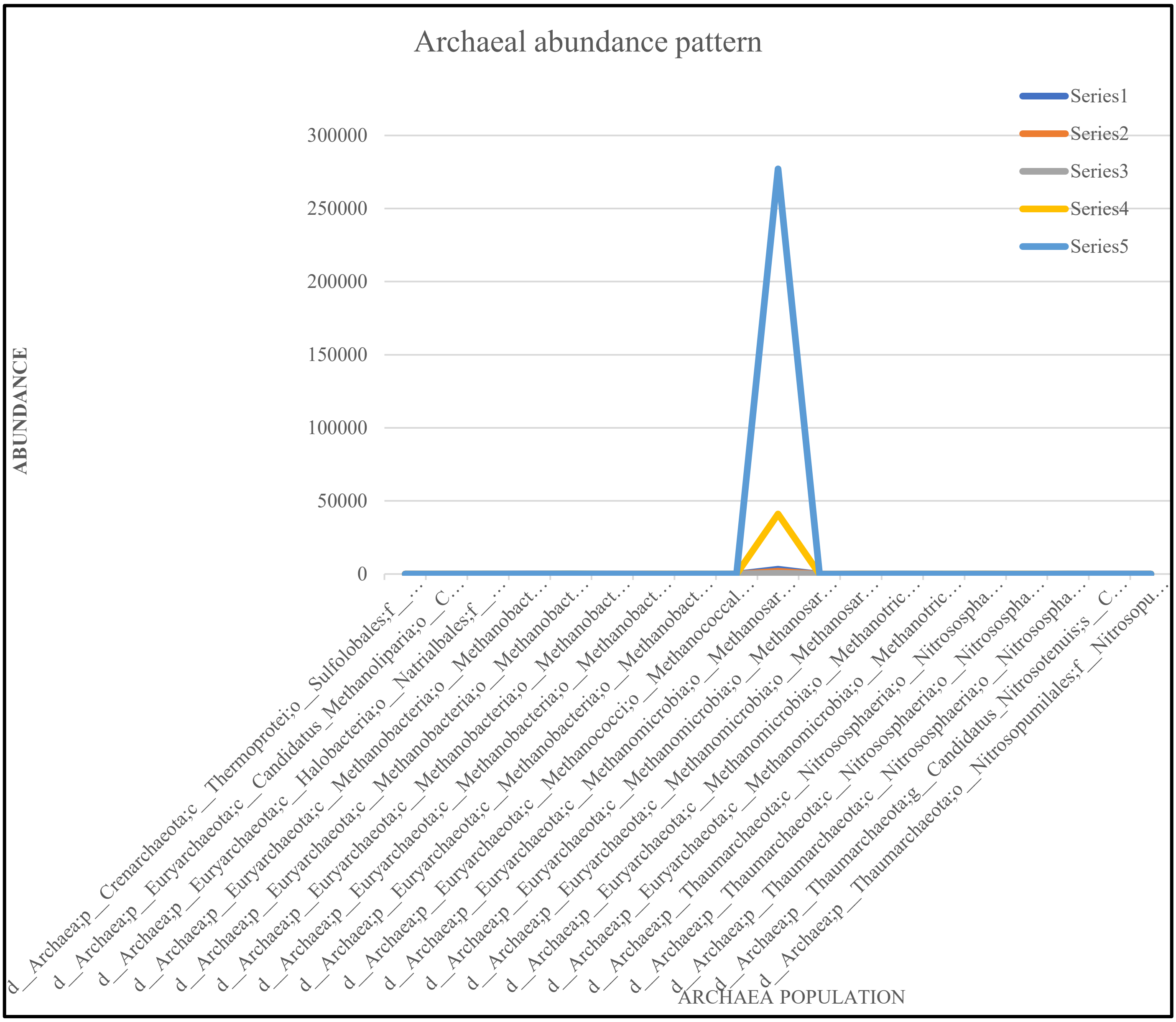


**Figure 6S** Archaeal abundance profile across different sites. Series 1, 2,3,4,5 relate the site 4, site 1, site 5, site 3 and site 2 respectively.


1. Bird, L. J.; Kuenen, J. G.; Osburn, M. R.; Tomioka, N.; Ishii, S.; Barr, C.; Nealson, K. H.; Suzuki, S., Serpentinimonas gen. nov., Serpentinimonas raichei sp. nov., Serpentinimonas barnesii sp. nov. and Serpentinimonas maccroryi sp. nov., hyperalkaliphilic and facultative autotrophic bacteria isolated from terrestrial serpentinizing springs. *Int. J. Syst. Evol. Microbiol.* **2021,** *71* (8).

2. Lee, E. M.; Jeon, C. O.; Choi, I.; Chang, K. S.; Kim, C. J., Silanimonas lenta gen. nov., sp. nov., a slightly thermophilic and alkaliphilic gammaproteobacterium isolated from a hot spring. *Int. J. Syst. Evol. Microbiol.* **2005,** *55* (Pt 1), 385-389.
3. Yan, Z.; Zhang, Y.; Wu, H.; Yang, M.; Zhang, H.; Hao, Z.; Jiang, H., Isolation and characterization of a bacterial strain Hydrogenophaga sp. PYR1 for anaerobic pyrene and benzo[a]pyrene biodegradation. *RSC Advances* **2017,** *7* (74), 46690-46698.
4. Kiessling, M.; Meyer, O., Profitable oxidation of carbon monoxide or hydrogen during heterotrophic growth of Pseudomonas carboxydoflava. *FEMS Microbiol. Lett.* **1982,** *13* (3), 333-338.
5. Choi, G. M.; Lee, S. Y.; Kim, S. Y.; Wee, J. H.; Im, W. T., Hydrogenophaga borbori sp. nov., isolated from activated sludge. *Int. J. Syst. Evol. Microbiol.* **2020,** *70* (1), 555-561.
6. Sayavedra, L.; Li, T.; Bueno Batista, M.; Seah, B. K. B.; Booth, C.; Zhai, Q.; Chen, W.; Narbad, A., Desulfovibrio diazotrophicus sp. nov., a sulfate-reducing bacterium from the human gut capable of nitrogen fixation. *Environ. Microbiol.* **2021,** *23* (6), 3164-3181.
7. Escalante-Semerena, J. C., Conversion of cobinamide into adenosylcobamide in bacteria and archaea. *J. Bacteriol.* **2007,** *189* (13), 4555-60.
8. de Crecy-Lagard, V.; Hutinet, G.; Cediel-Becerra, J. D. D.; Yuan, Y.; Zallot, R.; Chevrette, M. G.; Ratnayake, R.; Jaroch, M.; Quaiyum, S.; Bruner, S., Biosynthesis and function of 7-deazaguanine derivatives in bacteria and phages. *Microbiol. Mol. Biol. Rev.* **2024,** *88* (1), e0019923.
9. Fujita, S.; Sugio, Y.; Kawamura, T.; Yamagami, R.; Oka, N.; Hirata, A.; Yokogawa, T.; Hori, H., ArcS from <em>Thermococcus kodakarensis</em> transfers L-lysine to preQ<sub>0</sub> nucleoside derivatives as minimum substrate RNAs. *J. Biol. Chem.* **2024,** *300* (8).
